# Histone methyltransferase Ezh2 coordinates mammalian axon regeneration via epigenetic regulation of key regenerative pathways

**DOI:** 10.1101/2022.04.19.488817

**Authors:** Xue-Wei Wang, Shu-Guang Yang, Ming-Wen Hu, Rui-Ying Wang, Chi Zhang, Anish R. Kosanam, Arinze J. Ochuba, Jing-Jing Jiang, Ximei Luo, Jiang Qian, Chang-Mei Liu, Feng-Quan Zhou

**Author notes:** Correspondence to: Feng-Quan Zhou, Ph.D., Room 291, The John G. Rangos Sr. Building, 855 North Wolfe Street, Baltimore, MD 21205, USA., Chang-Mei Liu, Ph.D., Room 623, Institute for Stem Cell and Regeneration Building, 3 Datun Road, Chaoyang District, Beijing 100101, China. These authors contributed equally to this study (Xue-Wei Wang, Shu-Guang Yang, Ming-Wen Hu, Rui-Ying Wang).

## Abstract

Current clinical treatment for neurodegenerative diseases and neural injuries falls short of success, and one primary reason is that neurons in the mammalian central nervous system (CNS) lose their regeneration ability as they mature. Previous studies indicated that the regeneration ability of neurons is governed by complex signaling networks involving many genes. Therefore, here we investigated the roles of Ezh2, a histone methyltransferase, in regulation of mammalian axon regeneration at the epigenetic level. We found that Ezh2 level was gradually downregulated in the mouse nervous system during maturation but significantly upregulated in mature sensory neurons during spontaneous axon regeneration in the peripheral nerve system (PNS), suggesting its role in supporting axon regeneration. Indeed, *Ezh2* loss-of-function in sensory neurons impaired PNS axon regeneration *in vitro* and *in vivo*. In contrast, overexpression of *Ezh2* in retinal ganglion cells in the CNS induced optic nerve regeneration after optic nerve injury in both methyltransferase-dependent and -independent manners. Mechanistic exploration with multiomics sequencing, together with functional analyses, revealed that Ezh2 supported axon regeneration by systematically silencing the transcription of genes regulating synaptic function and axon regeneration inhibitory signaling, while broadly activating factors promoting axon regeneration. Our study not only reveals that Ezh2 coordinates axon regeneration via epigenetically regulating multiple key regenerative pathways, but also suggests that modulating chromatin accessibility is a promising strategy to promote CNS axon regeneration.

## Introduction

The common consequences of neurodegenerative diseases and neural injuries are axon degeneration and neuronal cell death. Failure of axon regeneration after injury or degeneration in the mammalian central nervous system (CNS) results in poor functional recovery and permanent disabilities. Unfortunately, current clinical therapeutics for neural injuries and neurodegenerative diseases still fall short of success. Therefore, understanding why mature neurons in the mammalian CNS cannot regrow axons has been a longstanding challenge in the field. Many studies over the past decades have revealed that the low intrinsic axon growth competency of mature CNS neurons (Curcio and Bradke, 2018; He and Jin, 2016; Mahar and Cavalli, 2018), together with extrinsic inhibitory molecules (Geoffroy and Zheng, 2014; Nogueira-Rodrigues et al., 2021; Schwab and Strittmatter, 2014), are the major contributors to the poor regeneration outcome. During development, young neurons are intrinsically competent in axon growth to establish neural circuits, whereas adult neurons possess poor axon growth ability to maintain the stability of the circuits. Moreover, the inhibitory extracellular environment also limits unnecessary axon sprouting, acting as another factor to stabilize the neural circuits (Yiu and He, 2006).

During maturation, the cellular state of neurons changes from favoring to limiting axon growth, likely regulated by modifications of the epigenomic and the subsequent transcriptomic landscape in neurons. Unlike CNS neurons, the axon regeneration ability of neurons in the peripheral nervous system (PNS) can be reactivated upon peripheral nerve injury by initiating a transcription-dependent regenerative response (Saijilafu et al., 2013; Smith and Skene, 1997). Recent studies have demonstrated that such a response also involves massive changes in the epigenome and transcriptome of PNS neurons (Palmisano et al., 2019; Renthal et al., 2020; Weng et al., 2016). Several studies also revealed that nerve injuries induced neural rejuvenation with a common developmental-like transcriptional program in sensory neurons (Chandran et al., 2016; Renthal *et al*., 2020; Zhou et al., 2006). Similar reversal to an embryonic transcriptomic state also occurs in mature corticospinal neurons at an early stage following spinal cord injury, although it cannot be sustained (Poplawski et al., 2020). Thus, it would be of great importance to reveal the epigenomic changes that occur during neuronal maturation or PNS neural regeneration. The knowledge gained could be used to epigenetically remodel the transcriptomic landscape of mature CNS neurons and allow them to reclaim their axon regeneration ability.

In this study, we investigated the role of an important epigenetic regulator, Ezh2 histone methyltransferase, in mammalian axon regeneration, because it is a major epigenetic regulator governing gene transcription. Ezh2 is the catalytic component of the polycomb repressive complex 2 (PRC2), which catalyzes the addition of methyl groups to lysine 27 on histone H3 (H3K27). Trimethylation of H3K27 (H3K27me3) is a heterochromatic histone modification, which condenses nearby chromatin to silence gene transcription (Margueron and Reinberg, 2011). Although most studies have focused on the role of Ezh2 as a histone methyltransferase, some studies clearly showed that under certain circumstances, Ezh2 could methylate non-histone substrates (Vasanthakumar et al., 2017; Xu et al., 2012) or exert methylation-independent functions (Kim et al., 2018; Lee et al., 2011; Shi et al., 2007), suggesting that it is a multifaceted molecule.

Here, we showed that the protein level of Ezh2 was downregulated in the nervous system during development and maturation and dramatically upregulated in PNS neurons during peripheral axon regeneration. Functionally, *Ezh2* knockdown or knockout in dorsal root ganglion (DRG) neurons impaired axon regeneration *in vitro* and *in vivo*. More importantly, overexpression of *Ezh2* in retinal ganglion cells (RGCs) significantly promoted optic nerve regeneration after an optic nerve crush (ONC) in both histone methylation-dependent and -independent manners. In addition, *Ezh2* overexpression also alleviated RGC death caused by the ONC or N-methyl-D-aspartate (NMDA)-induced excitotoxic retinal injury. Mechanistically, RNA-seq of enriched RGCs revealed that *Ezh2* overexpression decreased mRNA levels of many genes regulating neuronal excitability and synaptic transmission, most of which have important roles in mature RGCs. One of the most significantly downregulated genes was *Slc6a13*, which encodes GABA transporter 2, and its overexpression partially blocked optic nerve regeneration induced by *Ezh2* overexpression. In the meanwhile, Ezh2 also transcriptionally suppressed components of multiple classes of CNS axon regeneration inhibitory factors, including oligodendrocyte-myelin glycoprotein (OMgp), tenascin-R, Lingo3, and ephrin receptors EphA4/6/7/8. Overexpression of *Lingo3* or *Omg* (encoding OMgp) almost completely blocked the promoting effect of Ezh2 on optic nerve regeneration, indicating that Ezh2 functioned to relieve extrinsic inhibition on axon regeneration. Lastly, *Ezh2* overexpression induced upregulation of a group of well-known positive regulators of intrinsic axon regeneration capacity, such as c-Jun, c-Myc, Sox11, and osteopontin. Expression of *Wfdc1*, which has been shown to inhibit osteopontin expression (Ressler et al., 2014) and was also downregulated upon *Ezh2* overexpression, blocked Ezh2-induced optic nerve regeneration. Collectively, our results suggest that Ezh2 promotes mammalian axon regeneration through various molecular mechanisms, coordinately targeting both the intrinsic regenerative ability and extrinsic hostile environment.

## Results

### Ezh2 is developmentally downregulated in the nervous system and upregulated in adult sensory neurons after peripheral nerve injury

To evaluate how Ezh2 expression is regulated during neural development, we first examined the protein level of Ezh2 in mouse DRG and cortical tissues at different developmental stages. We found that Ezh2 was abundantly expressed in DRGs and the cortex in late embryonic stages, remained at high levels during the first several postnatal days, and then gradually declined to become hardly detectable at three to four weeks after birth (Fig. S1A, B). DRG neurons extend a single axon that bifurcates into two branches, a peripheral branch that readily regenerates upon injury in a transcription-dependent manner (Smith and Skene, 1997), and a central branch lacking the spontaneous regenerative ability. Sensory axons in the mouse sciatic nerve are primarily comprised of the peripheral branches of lumbar 4 and 5 (L4/5) DRG neurons. We observed a sharp increase of the protein level of Ezh2 in L4/5 DRGs three days after sciatic nerve transection (Fig. 1A). The results were consistent with a previous study showing increased Ezh2 and H3K27me3 levels in the DRG induced by spinal nerve ligation (Laumet et al., 2015), a neuropathic pain model that also injures the peripheral axons of DRG neurons. During development, sensory neurons gradually reduce their axon growth capacity after reaching their peripheral targets, correlating with the declined level of Ezh2. Conversely, the upregulation of Ezh2 after peripheral nerve injury accompanies the robust sensory axon regeneration, suggesting that Ezh2 might positively regulate axon regeneration. Similar changes of Ezh2 level in the cortex during development imply that Ezh2 might also be involved in regulation of axon growth and regeneration in the CNS.

**Figure 1.**
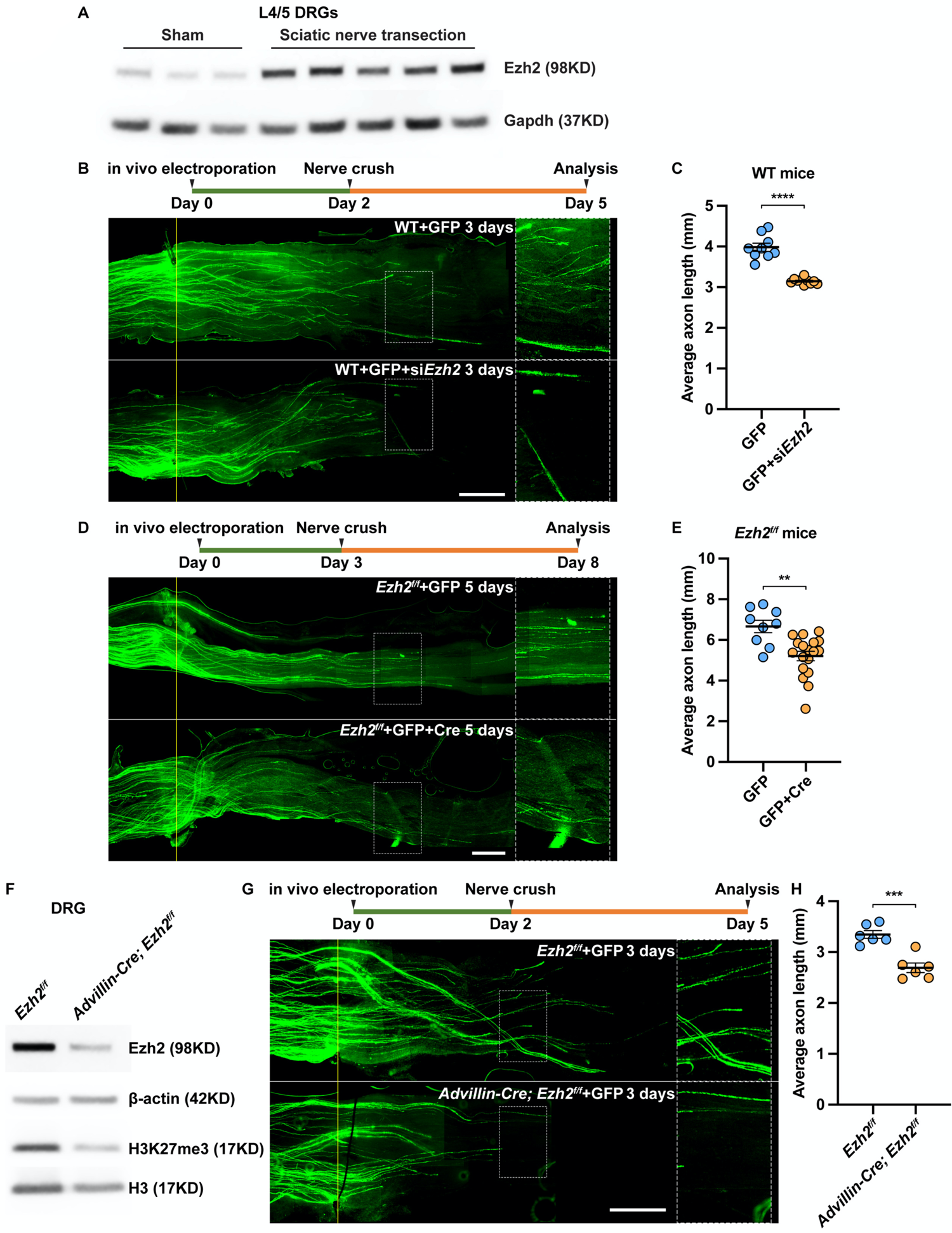
Upregulation of Ezh2 is necessary for spontaneous axon regeneration of sensory neurons *in vivo*. (A) Immunoblotting showing that Ezh2 is significantly increased in L4/5 DRGs 3 days after sciatic nerve transection (n = 3 for sham surgery, n = 5 for sciatic nerve transection). (B) Top: Time course of the experiment. Bottom: Representative images of sciatic nerves showing that *Ezh2* knockdown in L4/5 DRGs impairs spontaneous axon regeneration of sensory neurons *in vivo*. The right column displays enlarged images of the areas in white, dashed boxes on the left. The crush sites are aligned with the yellow line. Yellow arrows indicate longest axons in each nerve. Scale bar, 1 mm (0.5 mm for enlarged images). (C) Quantification of the average length of regenerating axons in (B) (unpaired t test; p < 0.0001; n = 9 nerves in the control condition, n = 10 nerves in the *Ezh2* knockdown condition). (D) Top: Time course of the experiment. Bottom: Representative images of sciatic nerves showing that *Ezh2* knockout in L4/5 DRGs impairs spontaneous axon regeneration of sensory neurons *in vivo*. The right column displays enlarged images of the areas in white, dashed boxes on the left. The crush sites are aligned with the yellow line. Yellow arrows indicate longest axons in each nerve. Scale bar, 1 mm (0.5 mm for enlarged images). (E) Quantification of the average length of regenerating axons in (D) (unpaired t test; p = 0.0011; n = 9 nerves in the control condition, n = 18 nerves in the *Ezh2* knockout condition). (F) Representative immunoblotting showing successful knockout of *Ezh2* and downregulation of H3K27me3 in DRG neurons of *Advillin-Cre; Ezh2^f/f^* mice (n = 3 independent experiments). (G) Top: Time course of the experiment. Bottom: Representative images of sciatic nerves showing that specific knockout of *Ezh2* in sensory neurons of L4/5 DRGs impairs spontaneous axon regeneration *in vivo*. The right column displays enlarged images of the areas in white, dashed boxes on the left. The crush sites are aligned with the yellow line. Yellow arrows indicate longest axons in each nerve. Scale bar, 1 mm (0.5 mm for enlarged images). (H) Quantification of the average length of regenerating axons in (G) (unpaired t test; p = 0.0003; n = 6 nerves in each condition). Data are represented as mean ± SEM. **p < 0.01, ***p < 0.001, ****p < 0.0001. See also Figure S1.

### Ezh2 is necessary for axon regeneration of sensory neurons *in vitro* and *in vivo*

To test our hypothesis, we first investigated if *Ezh2* loss-of-function would impair regenerative axon growth of cultured DRG neurons. Using *in vitro* electroporation (Zhou et al., 2004), siRNAs targeting *Ezh2* mRNA (siEzh2) were transfected into cultured DRG neurons. Control neurons were electroporated with non-targeting siRNAs (siNT). *Ezh2* was efficiently knocked down in the culture three days after the electroporation, which was confirmed by immunoblotting (Fig. S1C). Thus, on the 4^th^ day, we replated the neurons and cultured them for another 24 hours, as described in our earlier study (Saijilafu *et al*., 2013). The results showed that *Ezh2* knockdown significantly reduced regenerative axon growth by ∼25% within 24 hours (Fig. S1E, G). To rule out the possibility that the phenotype was caused by off-target effects of the siRNAs, we obtained *Ezh2^f/f^* mice and crossed them with *Advillin-Cre* mice (Zhou et al., 2010) to generate *Advillin-Cre; Ezh2^f/f^* mice, in which *Ezh2* was specifically deleted in sensory neurons. We performed a peripheral nerve conditioning lesion in *Advillin-Cre; Ezh2^f/f^* and *Ezh2^f/f^* (control) mice and waited for three days, and then cultured L4/5 DRG neurons for 24 hours. The results showed that *Ezh2* deletion significantly impaired regenerative axon growth of conditioning lesioned DRG neurons by ∼20% (Fig. S1F, H). Successful knockout of *Ezh2* was confirmed by immunoblotting of protein extracted from the cultured cells (Fig. S1D). The remaining Ezh2 signal most likely came from non-neuronal cells in the culture. These results demonstrated that Ezh2 supported regenerative axon growth of sensory neurons *in vitro*. To further explore if Ezh2 was also required for axon regeneration of DRG neurons *in vivo*, we knocked down *Ezh2* in L4/5 DRGs by *in vivo* electroporation of siEzh2, a technique widely used in our previous studies (Saijilafu et al., 2011). *CMV-GFP* plasmid was simultaneously electroporated to label the regenerating axons. Our previous study showed that *in vivo* electroporation of siNT had no effect on spontaneous axon regeneration of sensory neurons (Wang et al., 2018a). Therefore, control mice were electroporated with the *CMV-GFP* plasmid only. Two days after the electroporation, the sciatic nerves were crushed. Three days later, we found that *Ezh2* knockdown in DRGs significantly impaired sensory neuron axon regeneration *in vivo* by ∼20% (Fig. 1B, C). To rule out off-target effects of the siRNAs, we electroporated *CMV-Cre* and *CMV-GFP* plasmids into L4/5 DRGs of *Ezh2^f/f^* mice to knockout *Ezh2*. *Ezh2^f/f^* mice electroporated with the *CMV-GFP* plasmid only were used as the control group. To allow sufficient time for Cre-mediated recombination, the sciatic nerves were crushed three days after the electroporation. Five days after the sciatic nerve crush, we found that axon regeneration was significantly impaired by *Ezh2* knockout (Fig. 1D, E). To further rule out the possibility that the phenotype was caused by *Ezh2* loss-of-function in non-neuronal cells in the DRG, we electroporated the *CMV-GFP* plasmid in *Advillin-Cre; Ezh2^f/f^* and *Ezh2^f/f^* (control) mice and two days later, we crushed the sciatic nerves. Successful knockout of *Ezh2* and consequent downregulation of H3K27me3 in DRG neurons of *Advillin-Cre; Ezh2^f/f^* mice were confirmed by immunoblotting (Fig. 1F). Three days after the nerve crush, we found that specific deletion of *Ezh2* in sensory neurons significantly reduced axon regeneration *in vivo* by ∼20% (Fig. 1G, H). Collectively, these results provided clear evidence that Ezh2 was necessary for spontaneous axon regeneration of DRG neurons triggered by peripheral nerve injury both *in vitro* and *in vivo*.

### Ezh2 promotes optic nerve regeneration via histone methylation-dependent and -independent pathways

Since the injury-induced upregulation of Ezh2 is necessary for spontaneous axon regeneration of DRG neurons, we wondered if forced overexpression of *Ezh2* in RGCs could stimulate optic nerve regeneration. *Ezh2* was overexpressed in RGCs by intravitreal injection of AAV2-*Ezh2*. Control mice were injected with AAV2-*GFP*. Our previous studies showed that this approach could successfully transduce ∼90% of RGCs (Wang *et al*., 2018a; Wang et al., 2020). We confirmed successful overexpression of *Ezh2* at two weeks after the virus injection by immunoblotting of whole retinas (Fig. 2A) or RGCs enriched from dissociated retinal cells by fluorescence-activated cell sorting (FACS) (Fig. 2B). In addition, immunohistochemistry of retinal sections showed that H3K27me3 levels in RGCs were consequently increased (Fig. 2C, D). Two weeks after the virus injection, the optic nerves were crushed. *Ezh2* overexpression significantly improved the survival of RGCs two weeks after the ONC by ∼50% (Fig. 2E, F and Fig. S2). By using a different retinal injury model with intravitreal NMDA injection (Guo et al., 2021), we found that *Ezh2* overexpression almost doubled RGC survival one week after the excitotoxic injury to the retina (Fig. 2G, H), suggesting Ezh2 could protect RGCs from apoptosis independent of the retinal injury models. Optic nerve regeneration was also assessed two weeks after the ONC. The regenerating axons were labeled by Alexa Fluor-conjugated cholera toxin subunit B (CTB), which was intravitreally injected two days prior to tissue harvesting and anterogradely transported into the axons. Optic nerves were tissue-cleared and imaged with confocal microscopy. Compared to the control group, in which only a small number of axons were able to cross the crush site, overexpression of *Ezh2* induced markedly enhanced optic nerve regeneration after injury (Fig. 3A, C). Some axons grew longer than 1,250 μm in the two-week period (Fig. 3A, C).

**Figure 2.**
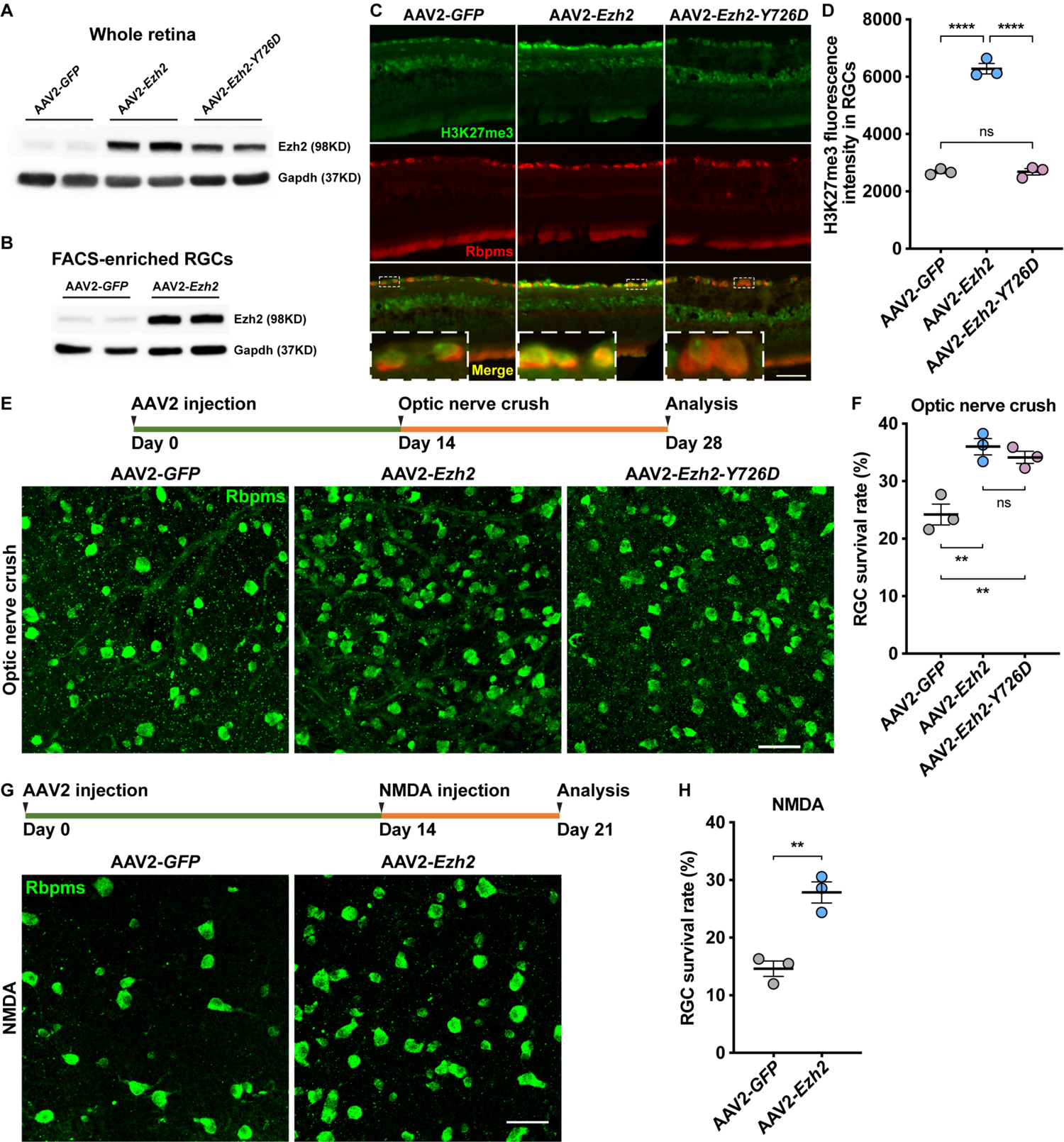
*Ezh2* overexpression enhances RGC survival after optic nerve crush or excitotoxic injury. (A, B) Immunoblotting showing increased Ezh2 level in the retina (A) or FACS-enriched RGCs (B) two weeks after intravitreal injection of AAV2-*Ezh2* or AAV2-*Ezh2-Y726D* (n = 2 in each condition). (C) Representative immunohistochemistry of retinal sections showing upregulated H3K27me3 levels in RGCs two weeks after intravitreal injection of AAV2-*Ezh2*, but not AAV2-*Ezh2-Y726D*. Retinal sections were stained with anti-H3K27me3 (green) and anti-Rbpms (red). Insets display enlarged images of RGCs in white, dashed boxes. Scale bar, 50 μm (12.5 μm for enlarged images). (D) Quantification of average fluorescence intensity of H3K27me3 immunoreactivity in RGCs in (C) (one-way ANOVA followed by Tukey’s multiple comparisons; p < 0.0001; n = 3 retinas in each condition; at least 150 RGCs from 10-12 non-adjacent sections were analyzed for each retina). (E) Top: Time course of the experiment. Bottom: Representative immunohistochemistry of whole-mount retinas showing that overexpression of *Ezh2* or *Ezh2-Y726D* improves RGC survival two weeks after optic nerve crush. Whole-mount retinas were stained with anti-Rbpms (green). Scale bar, 50 μm. (F) Quantification of RGC survival rate two weeks after optic nerve crush in (E) (one-way ANOVA followed by Tukey’s multiple comparisons; p = 0.0025; n = 3 pairs of retinas in each condition; 6-9 fields were analyzed for each retina). (G) Top: Time course of the experiment. Bottom: Representative immunohistochemistry of whole-mount retinas showing that overexpression of *Ezh2* improves RGC survival one week after NMDA-induced excitotoxic injury. Whole-mount retinas were stained with anti-Rbpms (green). Scale bar, 50 μm. (H) Quantification of RGC survival rate one week after NMDA-induced excitotoxic injury in (F) (unpaired t test; p = 0.0042; n = 3 pairs of retinas in each condition; 6-8 fields were analyzed for each retina). Data are represented as mean ± SEM. *P values* of post hoc analyses are illustrated in the figure. ns, not significant, **p < 0.01, ****p < 0.0001. See also Figure S2.

**Figure 3.**
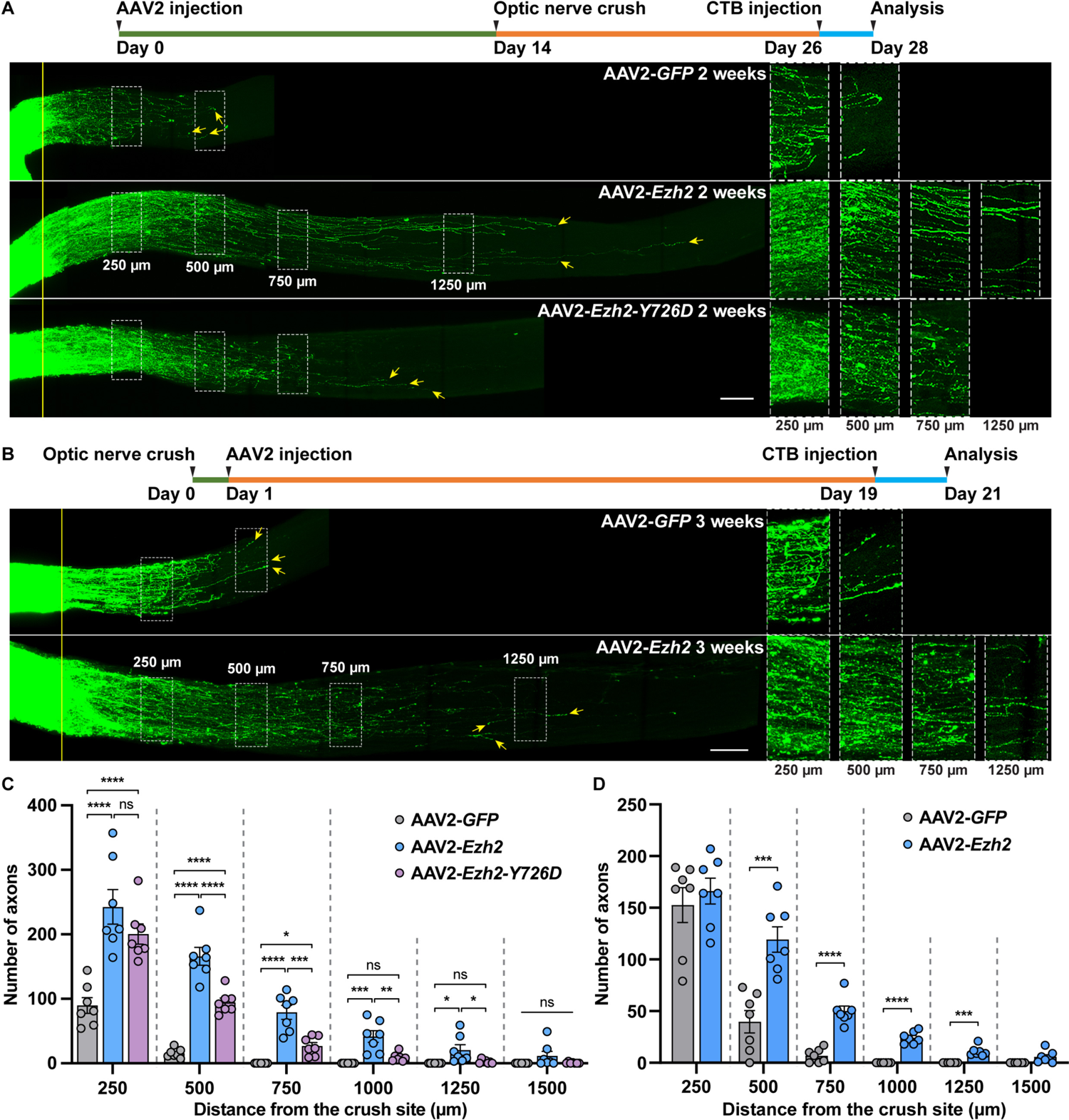
Ezh2 supports optic nerve regeneration via both methyltransferase activity-dependent and -independent pathways. (A) Top: Time course of the experiment. Bottom: Representative images of optic nerves showing that pre-injury overexpression of *Ezh2* induces strong optic nerve regeneration two weeks after optic nerve crush. Overexpression of *Ezh2-Y726D* also modestly promotes optic nerve regeneration. Columns on the right display enlarged images of the areas in white, dashed boxes on the left, showing axons at 250, 500, 750, and 1250 μm distal to the crush sites, which are aligned with the yellow line. Yellow arrows indicate longest axons in each nerve. Scale bar, 100 μm (50 μm for enlarged images). (B) Top: Time course of the experiment. Bottom: Representative images of optic nerves showing that post-injury overexpression of *Ezh2* induces significant optic nerve regeneration. Columns on the right display enlarged images of the areas in white, dashed boxes on the left, showing axons at 250, 500, 750, and 1250 μm distal to the crush sites, which are aligned with the yellow line. Yellow arrows indicate longest axons in each nerve. Scale bar, 100 μm (50 μm for enlarged images). (C) Quantification of optic nerve regeneration in (A) (one-way ANOVA followed by Tukey’s multiple comparisons; p < 0.0001 at 250, 500, 750, and 1,000 μm; p = 0.0126 and 0.1029 at 1,250 and 1,500 μm, respectively; n = 7 nerves in each condition). (D) Quantification of optic nerve regeneration in (B) (unpaired t test; p = 0.5305, 0.0004, 0.0003, and 0.0545 at 250, 500, 1,250, and 1,500 μm, respectively; p < 0.0001 at 750 and 1,000 μm; n = 7 nerves in each condition). Data are represented as mean ± SEM. *P values* of post hoc analyses are illustrated in the figure. ns, not significant, *p < 0.05, **p < 0.01, ***p < 0.001, ****p < 0.0001.

To investigate if the histone methyltransferase activity of Ezh2 was required for its role in promoting optic nerve regeneration and RGC survival, we constructed a vector expressing a mutant form of Ezh2, whose 726^th^ amino acid was mutated from a tyrosine to an aspartic acid (Ezh2-Y726D), and overexpressed it in RGCs (Fig. 2A). Previous studies reported that this single amino acid mutation could eliminate the methyltransferase activity of both human and mouse Ezh2 (Ernst et al., 2010; Lavarone et al., 2019). This was confirmed by our immunohistochemistry results of retinal sections, in which overexpression of *Ezh2-Y726D* had no effect on H3K27me3 levels in RGCs at all (Fig. 2C, D). To our surprise, this catalytically dead Ezh2 mutant promoted RGC survival to the same extent as did wild type Ezh2 two weeks after the ONC (Fig. 2E, F and Fig. S2), suggesting that its function in preventing cell death was histone methylation independent. Moreover, Ezh2-Y726D also induced optic nerve regeneration, although to a much lesser extent (Fig. 3A, C), indicating that both histone methylation-dependent and -independent mechanisms contributed to the role of Ezh2 in promoting axon regeneration.

To explore the translational potential of Ezh2 in CNS neural regeneration, we tested if post-injury overexpression of *Ezh2* in RGCs could also promote optic nerve regeneration. We first performed the ONC and one day later, AAV2-*GFP* or AAV2-*Ezh2* was intravitreally injected. Optic nerve regeneration was examined three weeks after the ONC. We found that post-injury *Ezh2* overexpression still overtly promoted optic nerve regeneration compared to that in the control condition (Fig. 3B, D), despite it was weaker than that induced by pre-injury overexpression of *Ezh2*. Specifically, more regenerating axons could be found from 500 to 1,250 μm distal of the crush site after Ezh2 overexpression (Fig. 3B, D). However, equivalent numbers of regenerating axons were observed at 250 μm from the crush site between the two conditions (Fig. 3B, D). This was likely caused by the extended regeneration period in the control group (three weeks in Fig. 3B, D vs. two weeks in Fig. 3A, C) and delayed *Ezh2* expression in RGCs. These results demonstrated the translational potential of *Ezh2* gain-of-function for enhancing axon regeneration in the CNS.

### Ezh2 modifies the RGC transcriptome to regulate multiple categories of target genes

To gain mechanistic insights into how Ezh2 supports RGC axon regeneration, we profiled the transcriptomic and epigenomic changes in RGCs induced by *Ezh2* or *Ezh2-Y726D* overexpression with RNA sequencing (RNA-seq) and assay for transposase-accessible chromatin with sequencing (ATAC-seq). We intravitreally injected AAV2-*GFP* (control), AAV2-*Ezh2*, or AAV2-*Ezh2-Y726D*, allowed two weeks for efficient transgene expression, and crushed the optic nerves. Three days after the ONC, RGCs were enriched from dissociated retinal cells with FACS, and RNA-seq or ATAC-seq libraries were constructed (Fig. S3A-D and Fig. S4A-C). One RNA-seq library from the *Ezh2* overexpression condition was excluded from data analysis based on results of principle component analysis (PCA) and hierarchical clustering analysis (Fig. S3A, B). Chromatin accessibility at the promoter region moderately correlated with RNA expression within each condition (Fig. S4D-G), suggesting consistency between the RNA-seq and ATAC-seq.

We identified 669 differentially expressed genes (DEGs) in the RNA-seq between control and *Ezh2* overexpression conditions at the threshold of adjusted *P* value < 0.05 and fold change > 1.5 (Fig. 4A, Fig. S3E, and Table S1). Surprisingly, despite being a catalytically dead form of *Ezh2*, *Ezh2-Y726D* overexpression resulted in 1,103 DEGs (Fig. 4B and Table S1), which was a lot more than those induced by *Ezh2* overexpression. This was consistent with our ATAC-seq results, in which considerably more differentially accessible regions were found after *Ezh2-Y726D* overexpression (Table S2), suggesting that unknown transcriptional regulation activities of Ezh2 independent of the methyltransferase function are yet to be discovered. We then examined how the 669 DEGs induced by *Ezh2* overexpression are regulated by *Ezh2-Y726D* overexpression. Although not all of them were significantly regulated by Ezh2-Y726D, the heatmaps indicated that most of them showed opposite patterns of regulation after *Ezh2* or *Ezh2-Y726D* overexpression (Fig. S3E). Indeed, among the 236 common DEGs regulated by both Ezh2 and Ezh2-Y726D, 203 (86%) changed in opposite directions (Fig. S3F and Table S1). Gene ontology (GO) analysis of the DEGs further revealed that Ezh2 and Ezh2-Y726D inversely modified the RGC transcriptome (Fig. 4C and Table S3). Specifically, *Ezh2* overexpression led to downregulation of a series of ion transport and synaptic transmission-related genes and upregulation of many immune response programs, both of which were oppositely regulated by Ezh2-Y726D overexpression (Fig. 4C-E). Similarly, a large number of GO terms (immune response genes) were shared by genes whose promoter regions became more open after *Ezh2* overexpression and those whose promoter regions became more closed after *Ezh2-Y726D* overexpression in the ATAC-seq (Fig. S4H and Table S4). These results implied that Ezh2-Y726D might act to inhibit functions of endogenous Ezh2 in a dominant negative manner.

**Figure 4.**
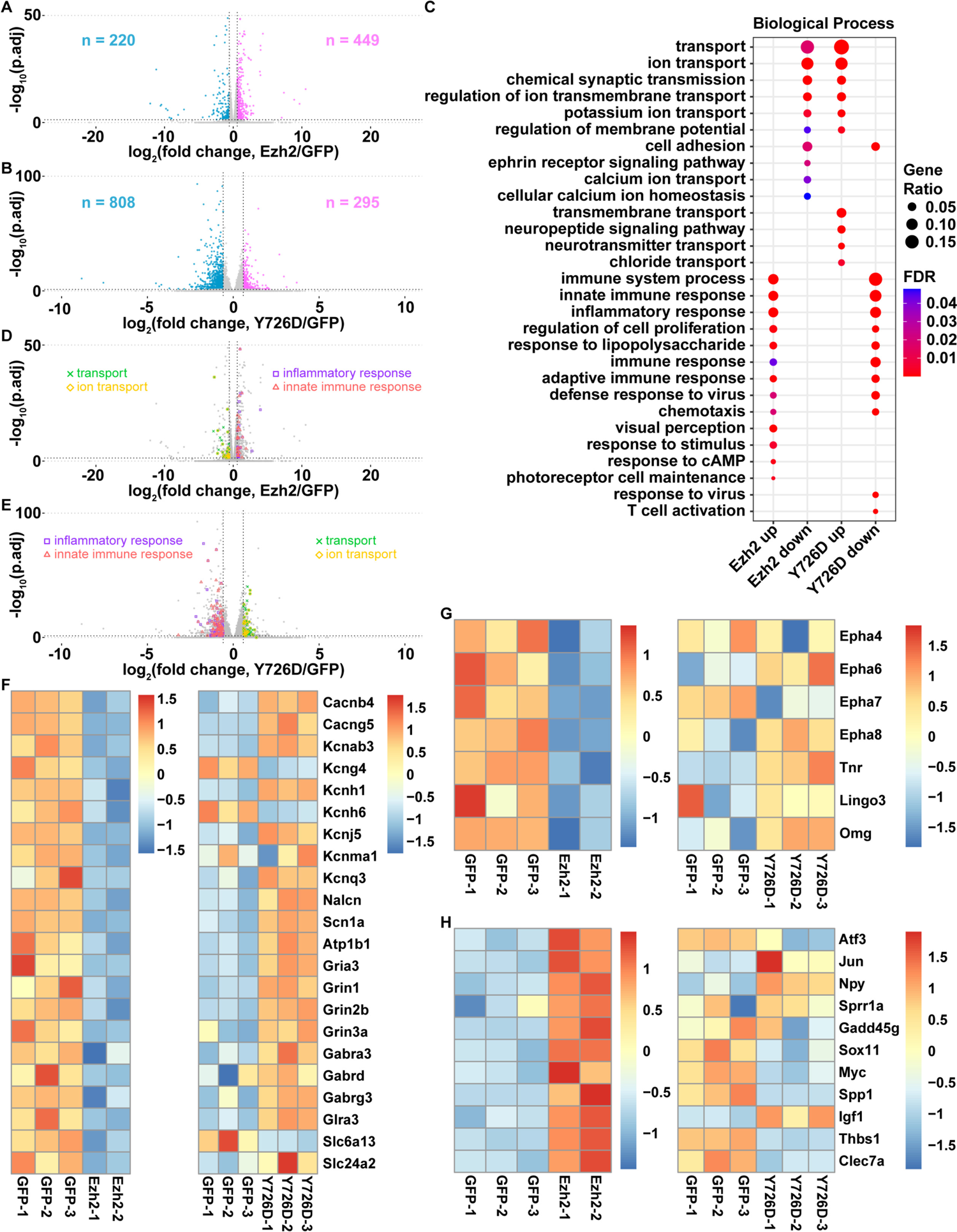
Ezh2 modifies the RGC transcriptome to regulate multiple categories of target genes. (A, B) Volcano plots showing differences in gene expression between control and *Ezh2* overexpression conditions (A) or between control and *Ezh2-Y726D* overexpression conditions (B). Note that 12 genes with -log_10_(p.adj) > 50 and 3 genes with -log_10_(p.adj) > 100 are not plotted in (A) and (B), respectively. (C) Gene ontology (GO) analysis of differentially expressed genes (DEGs) induced by *Ezh2* or *Ezh2-Y726D* overexpression. A subset of most significantly enriched GO terms in the biological process category are shown here. (D, E) Volcano plots described in (A, B) with DEGs in four enriched GO terms labeled. (F, G) Heatmaps of mRNA levels of neuronal excitability and synaptic transmission regulators (F) or axon regeneration inhibitory factors (G) downregulated by *Ezh2* overexpression in the control vs. *Ezh2* overexpression RNA-seq (left) or the control vs. *Ezh2-Y726D* overexpression RNA-seq (right). (H) Heatmaps of mRNA levels of axon regeneration positive regulators upregulated by *Ezh2* overexpression in the control vs. *Ezh2* overexpression RNA-seq (left) or the control vs. *Ezh2-Y726D* overexpression RNA-seq (right). Note that the control vs. *Ezh2* overexpression RNA-seq and the control vs. *Ezh2-Y726D* overexpression RNA-seq were performed independently, therefore control (GFP) libraries in one experiment were independent of those in the other one. See also Figure S3-S5.

Because Ezh2 primarily functions to repress gene transcription through H3K27me3, we first focused on genes that were downregulated by *Ezh2* overexpression. GO analysis showed that *Ezh2* overexpression decreased mRNA levels of many genes encoding ion channels and transporters as well as neurotransmitter receptors and transporters (Fig. 4C, D, F), which are all important regulators of neuronal excitability and synaptic transmission. Except for a few genes, most of them were upregulated by Ezh2-Y726D (Fig. 4F), indicating that the regulation was mainly H3K27me3 dependent. Since neuronal excitability and synaptic transmission are fundamental biological functions of mature neurons, these results suggested that Ezh2 might promote axon regeneration by rejuvenating mature CNS neurons at the transcriptomic level.

Close examination of the downregulated genes further revealed that Ezh2 also reduced mRNA levels of multiple axon regeneration inhibitory factors or their receptors, including ephrin receptors (encoded by *Epha4, 6, 7, 8*; note that *Ehpa4* was not among the 669 DEGs but had an adjusted *P* value < 0.05), tenascin-R (encoded by *Tnr*), Lingo3, and OMgp (encoded by *Omg*) (Fig. 4G). By using an available single-cell RNA-seq dataset of adult mouse RGCs (https://singlecell.broadinstitute.org/single_cell/study/SCP509/mouse-retinal-ganglion-cell-adult-atlas-and-optic-nerve-crush-time-series) (Tran et al., 2019), we confirmed that all these inhibitory molecules or receptors were expressed in most RGCs (Fig. S5A-G). Moreover, the transcription of these genes was not repressed by Ezh2-Y726D (Fig. 4G), suggesting the involvement of H3K27me3 in regulation of these regeneration inhibitors by Ezh2.

In addition to suppressing gene transcription, *Ezh2* overexpression also induced upregulation of many well-known positive regulators of axon regeneration (Fig. 4H). Among them, *Atf3*, *Jun*, *Npy*, *Sprr1a*, *Gadd45g*, and *Sox11* are some of the first identified regeneration-associated genes (Befort et al., 2003; Bonilla et al., 2002; Jenkins and Hunt, 1991; Tanabe et al., 2003; Tsujino et al., 2000; Wakisaka et al., 1991). Others, including *Myc*, *Spp1* (encoding osteopontin), *Igf1*, *Thbs1* (encoding thrombospondin-1), and *Clec7a* (encoding dectin-1), have been well documented to promote CNS axon regeneration (Baldwin et al., 2015; Belin et al., 2015; Bray et al., 2019; Duan et al., 2015; Qian and Zhou, 2020; Yang et al., 2020). In contrast, *Ezh2-Y726D* overexpression downregulated many of these genes (Fig. 4H), indicating that their upregulation after *Ezh2* overexpression was mostly H3K27me3 dependent. Since H3K27me3 is a heterochromatic histone modification, the increase of these positive regulators of axon regeneration was most likely secondary to the elevation of H3K27me3.

Interestingly, Ezh2-Y726D also regulated the transcription of some genes in the same direction as did wild type Ezh2, including *Kcng4*, *Kcnh6*, and *Slc6a13* in the ion and neurotransmitter transport group (Fig. 4F), as well as *Jun*, *Npy*, and *Igf1* in the axon regeneration enhancer group (Fig. 4H). We think that these changes induced by Ezh2-Y726D might explain its weak promoting effect on optic nerve regeneration (see Fig. 3A, C).

### Ezh2 supports optic nerve regeneration by downregulating synaptic function-related genes

To determine if genes regulated by Ezh2 functionally act downstream to regulate axon regeneration, we first examined the function of *Slc6a13*, which encodes GABA transporter 2 and was one of the genes most significantly downregulated by *Ezh2* overexpression (see Fig. 4F). Using the available single-cell RNA-seq dataset of adult mouse RGCs (Tran *et al*., 2019), we verified that *Slc6a13* was broadly expressed by RGC subtypes (Fig. S5H). Additionally, by using the cleavage under targets and tagmentation (CUT&Tag) method (Kaya-Okur et al., 2019) followed by quantitative polymerase chain reaction (qPCR), we found that the promoter region of *Slc6a13* was bound by H3K27me3 (Fig. 5A), indicating *Slc6a13* transcription can be directly regulated by Ezh2 via H3K27me3. Functionally, when *Slc6a13* was overexpressed along with *Ezh2* in RGCs by intravitreal injection of AAV2-*Slc6a13*, the strong optic nerve regeneration stimulated by *Ezh2* overexpression was largely blocked, whereas *Slc6a13* overexpression *per se* did not have any effect (Fig. 5A, B). These results demonstrated that Ezh2 enhanced optic nerve regeneration, at least in part, by repressing *Slc6a13* transcription, and suggested that extracellular GABA level might play an important role in regulation of optic nerve regeneration. A previous study in larval sea lampreys showed that GABA promoted survival and axon regeneration of descending neurons after a complete spinal cord injury through GABA_B_ receptors (Romaus-Sanjurjo et al., 2018). Two recent studies showed that activating GABA_B_ receptors could promote axon regeneration in the mouse dorsal column and cortical spinal tract after spinal cord injury (Hilton et al., 2022; Li et al., 2020). We also investigated the role of *Slc6a13* in regenerative axon growth of DRG neurons *in vitro*. Consistently, we found that *Slc6a13* overexpression in sensory neurons had a mild but significant inhibitory effect on axon regeneration (Fig. S6A, B), suggesting its broad role in controlling axon regeneration.

**Figure 5.**
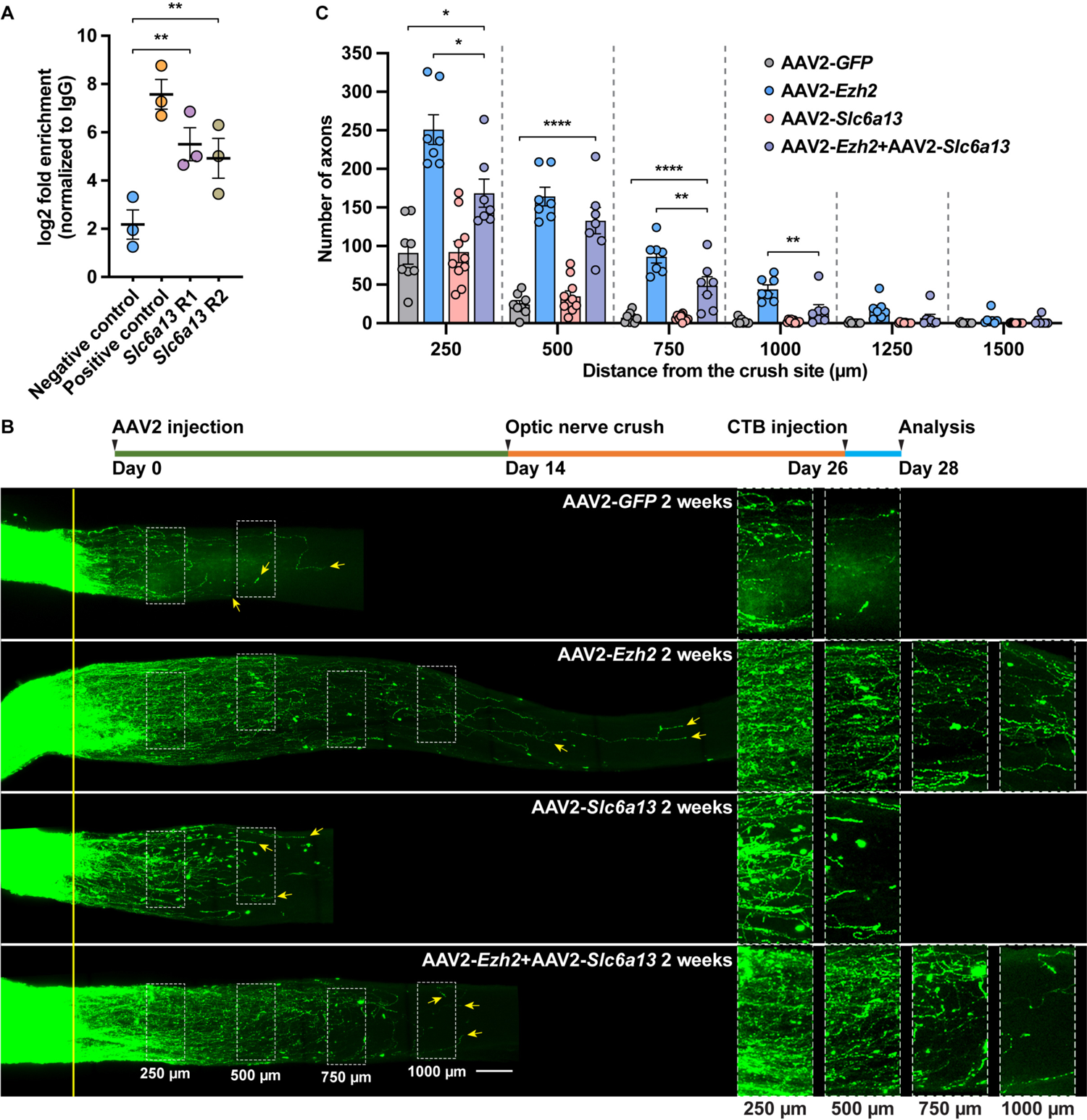
Ezh2 supports optic nerve regeneration by transcriptionally suppressing *Slc6a13*. (A) CUT&Tag followed by qPCR analysis showing that H3K27me3 is highly enriched in the promoter region of *Slc6a13* (paired t test; p = 0.0019 between negative control and *Slc6a13* R1; p = 0.0099 between negative control and *Slc6a13* R2; n = 3 independent experiments). (B) Top: Time course of the experiment. Bottom: Representative images of optic nerves showing that *Slc6a13* overexpression partially blocks *Ezh2* overexpression induced optic nerve regeneration. Columns on the right display enlarged images of the areas in white, dashed boxes on the left, showing axons at 250, 500, 750, and 1000 μm distal to the crush sites, which are aligned with the yellow line. Yellow arrows indicate longest axons in each nerve. Scale bar, 100 μm (50 μm for enlarged images). (C) Quantification of optic nerve regeneration in (A) (one-way ANOVA followed by Tukey’s multiple comparisons; p < 0.0001 at 250, 500, 750, and 1,000 μm; p = 0.0010 and 0.1099 at 1,250 and 1,500 μm, respectively; n = 8 nerves in the control condition, n = 10 nerves in the *Slc6a13* overexpression condition, n = 7 nerves in other conditions). Data are represented as mean ± SEM. *P values* of post hoc analyses are illustrated in the figure. *p < 0.05, **p < 0.01, ****p < 0.0001. See also Figure S5, S6.

### Ezh2 enhances optic nerve regeneration by suppressing major axon regeneration inhibitory signaling

It is widely accepted that CNS axon regeneration inhibitors can be categorized into three classes, which are MAIs, CSPGs, and repulsive axon guidance cues (Giger et al., 2010). Our results showed that Ezh2 silenced transcription of regeneration inhibitors or their receptors associated with all three classes (see Fig. 4G), indicating that Ezh2 might be a master suppressor of CNS axon regeneration inhibitory signaling. We thus tested if downregulation of *Omg* and *Lingo3* contributed to *Ezh2* overexpression-induced optic nerve regeneration. Overexpression of *Omg* or *Lingo3 per se* did not affect optic nerve regeneration, but almost completely blocked *Ezh2* overexpression-induced regeneration. Only at 250 μm from the crush site did we observe more regenerating axons in the co-overexpression groups than in the control group (Fig. 6A, B). Similarly, CUT&Tag followed by qPCR revealed that the *Lingo3* promoter region was bound by H3K27me3 (Fig. 6C), suggesting H3K27me3-mediated transcriptional repression of *Lingo3* following *Ezh2* overexpression. In contrast, no enrichment of the *Omg* promoter region was detected using an H3K27me3 antibody compared to normal IgG (Fig. 6C), suggesting that *Omg* downregulation might be a secondary effect of H3K27me3 upregulation. Collectively, these results demonstrated that Ezh2 supported optic nerve regeneration by transcriptionally silencing major axon regeneration inhibitory signaling and provided clear evidence that neurons could inhibit their own axon regeneration in an autocrine manner.

**Figure 6.**
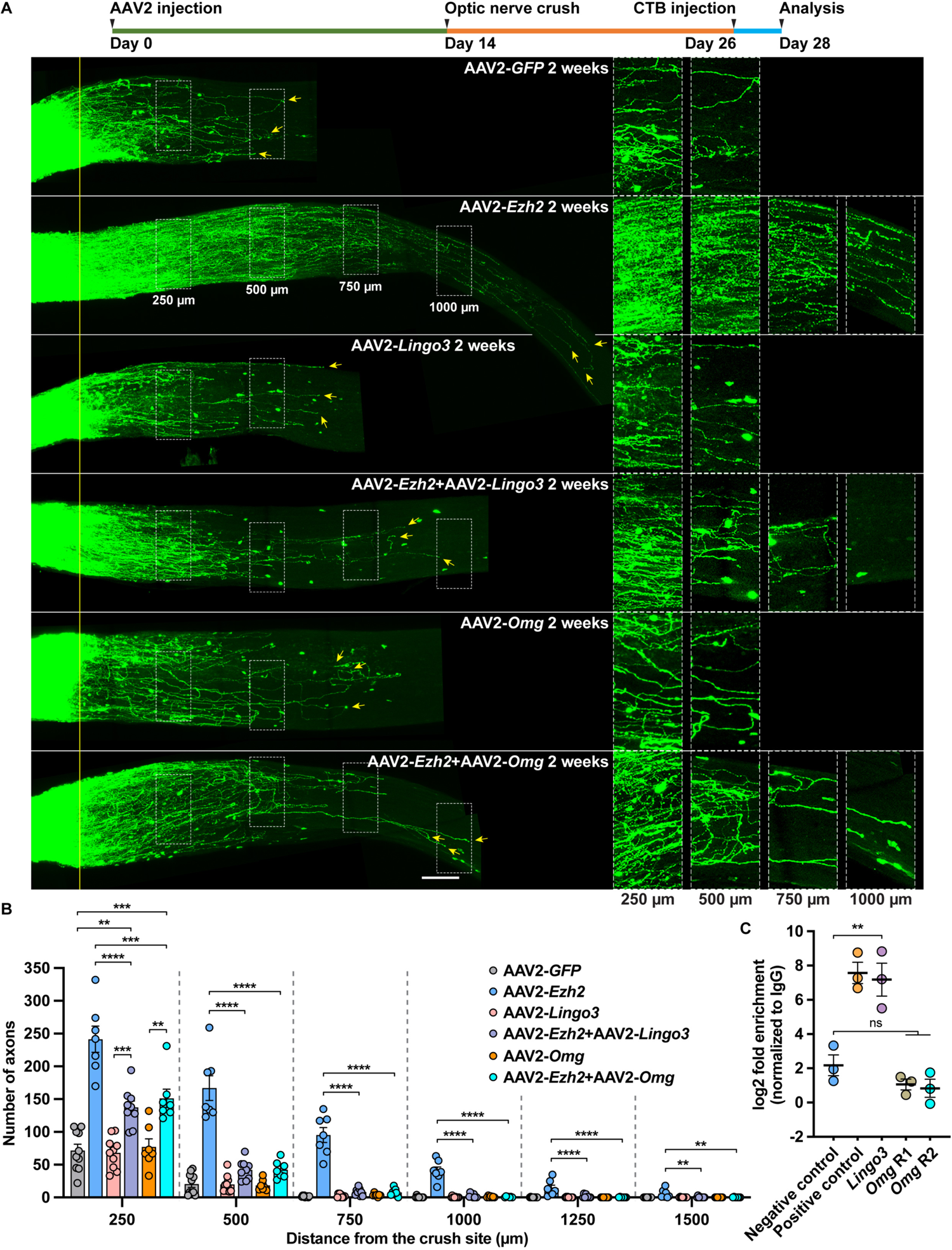
Ezh2 supports optic nerve regeneration by transcriptionally suppressing *Lingo3* and *Omg*. (A) Top: Time course of the experiment. Bottom: Representative images of optic nerves showing that *Lingo3* or *Omg* overexpression almost completely blocks *Ezh2* overexpression induced optic nerve regeneration. Columns on the right display enlarged images of the areas in white, dashed boxes on the left, showing axons at 250, 500, 750, and 1000 μm distal to the crush sites, which are aligned with the yellow line. Yellow arrows indicate longest axons in each nerve. Scale bar, 100 μm (50 μm for enlarged images). (B) Quantification of optic nerve regeneration in (A) (one-way ANOVA followed by Tukey’s multiple comparisons; p < 0.0001 at 250, 500, 750, 1,000, and 1,250 μm; p = 0.0003 at 1,500 μm; n = 10 nerves in the control condition and the *Lingo3* overexpression condition, n = 9 nerves in the *Ezh2* and *Lingo3* co-overexpression condition, n = 7 nerves in other conditions). (C) CUT&Tag followed by qPCR analysis showing that H3K27me3 is highly enriched in the promoter region of *Lingo3*, but not that of *Omg* (paired t test; p = 0.0058 between negative control and *Lingo3* R1; p = 0.1528 between negative control and *Omg* R1; p = 0.1815 between negative control and *Omg* R2; n = 3 independent experiments). Note that negative control and positive control are identical to those in Figure 5A. Data are represented as mean ± SEM. *P values* of post hoc analyses are illustrated in the figure. ns, not significant, **p < 0.01, ***p < 0.001, ****p < 0.0001. See also Figure S5.

### Ezh2 activates multiple axon regeneration enhancing pathways

Ezh2 overexpression resulted in upregulation of osteopontin (encoded by *Spp1*) in RGCs (see Fig. 4H and Table S1), which was reported to selectively promote αRGC axon regeneration (Duan *et al*., 2015). In addition to retinal repair, increased osteopontin expression has also been shown to underlie the enhanced tissue repair induced by knocking out *Wfdc1* (Ressler *et al*., 2014), a gene known as a tumor suppressor (Zhu et al., 2021) and a wound repair inhibitor (Ressler *et al*., 2014). Interestingly, the mRNA level of *Wfdc1* in RGCs was also reduced by Ezh2 in our RNA-seq results (Table S1), suggesting that it might regulate Ezh2-induced optic nerve regeneration via osteopontin. Indeed, overexpression of *Wfdc1* alone had little effect, whereas it completely blocked the optic nerve regeneration induced by *Ezh2* overexpression (Fig. 7A, B), suggesting that *Wfdc1* downregulation might be necessary for Ezh2-induced optic nerve regeneration. CUT&Tag followed by qPCR showed that the *Wfdc1* promoter region was bound by H3K27me3 (Fig. 7C), indicating Ezh2 could directly repress its transcription via its histone methyltransferase activity. These results provided a potential mechanism of how Ezh2 indirectly upregulates axon regeneration enhancing factors.

**Figure 7.**
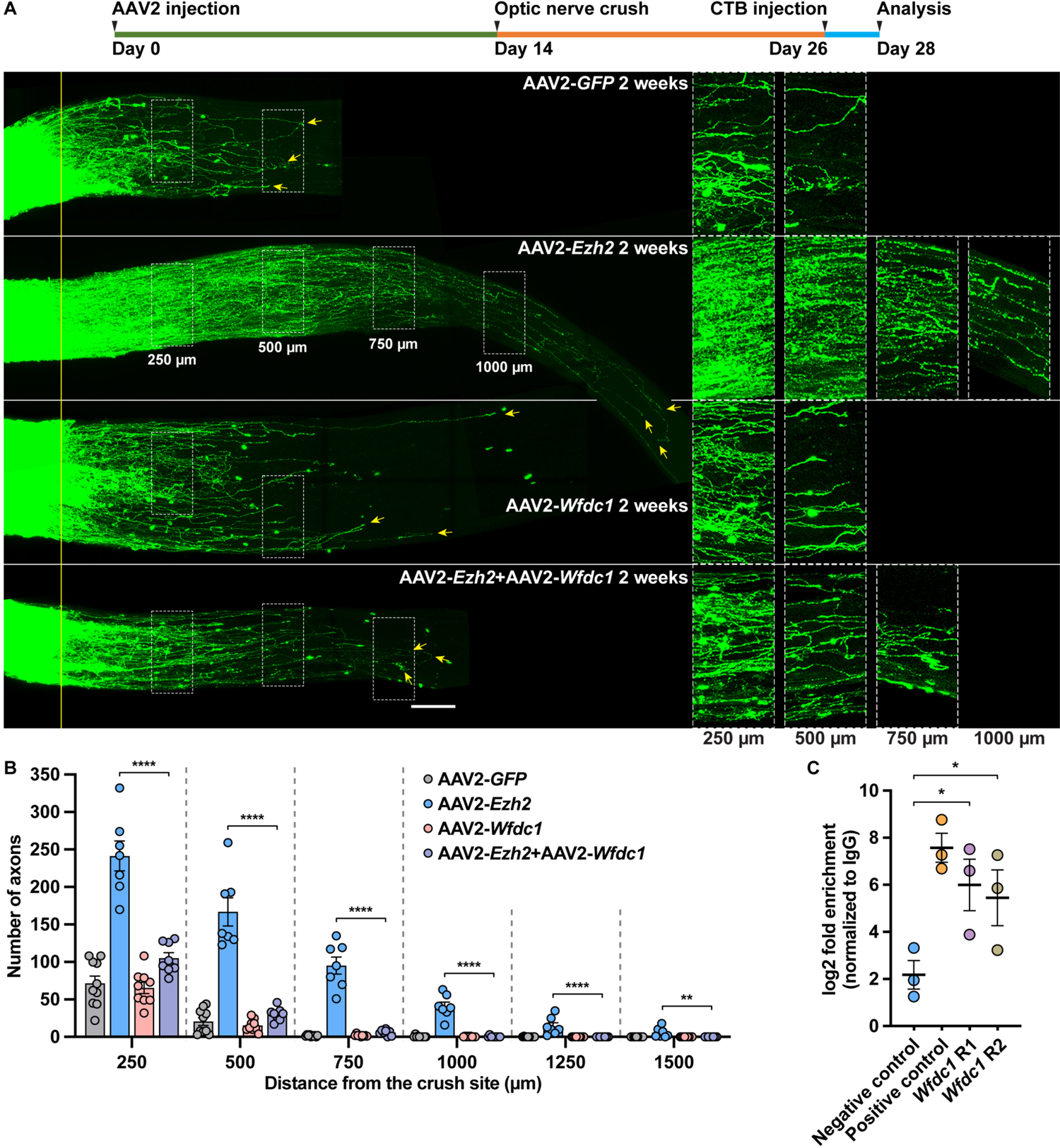
Ezh2 supports optic nerve regeneration by transcriptionally suppressing *Wfdc1*. (A) Top: Time course of the experiment. Bottom: Representative images of optic nerves showing that *Wfdc1* overexpression completely blocks *Ezh2* overexpression induced optic nerve regeneration. Columns on the right display enlarged images of the areas in white, dashed boxes on the left, showing axons at 250, 500, 750, and 1000 μm distal to the crush sites, which are aligned with the yellow line. Yellow arrows indicate longest axons in each nerve. Scale bar, 100 μm (50 μm for enlarged images). (B) Quantification of optic nerve regeneration in (A) (one-way ANOVA followed by Tukey’s multiple comparisons; p < 0.0001 at 250, 500, 750, 1,000, and 1,250 μm; p = 0.0015 at 1,500 μm; n = 10 nerves in the control condition, n = 7 in the *Ezh2* overexpression condition, n = 9 in the *Wfdc1* overexpression condition, n = 8 in the *Ezh2* and *Wfdc1* co-overexpression condition). Note that the control and *Ezh2* overexpression groups are identical to those in Figure 6B. (C) CUT&Tag followed by qPCR analysis showing that H3K27me3 is highly enriched in the promoter region of *Wfdc1* (paired t test; p = 0.0248 between negative control and *Wfdc1* R1; p = 0.0375 between negative control and *Wfdc1* R2; n = 3 independent experiments). Note that negative control and positive control are identical to those in Figure 5A. Data are represented as mean ± SEM. *P values* of post hoc analyses are illustrated in the figure. *p < 0.05, **p < 0.01, ****p < 0.0001. See also Figure S7.

Among other upregulated genes (Table S1), *Ascl1* and *Neurog2* (encoding Neurog2, also known as neurogenin-2) are well known to play important roles in regulation of neurogenesis and axon guidance during development (Dennis et al., 2019). They have also been shown to be direct reprogramming factors converting glial cells into neurons (Dennis *et al*., 2019). Moreover, Ascl1, Neurog2 and Ezh2 have recently been identified to be key factors driving neuronal differentiation in a Crispr-based screening study (Liu et al., 2018), suggesting similar functions between these genes. Indeed, Ascl1 has been shown to support PNS axon regeneration in mice (Lisi et al., 2017) and CNS axon regeneration in zebrafish and rats (Williams et al., 2015). We therefore investigated if Neurog2 could also regulate optic nerve regeneration. To our surprise, the results showed that overexpression of *Neurog2* in RGCs had little effect on optic nerve regeneration (Fig. S7A, B), suggesting distinct mechanisms mediating neurogenesis and axon regeneration.

### Ezh2 overexpression does not alter the epigenetic aging clock of RGCs

A recent study discovered that ONC increased the DNA methylation age of RGCs, and polycistronic expression of reprogramming factors Oct4, Sox2, and Klf4 counteracted the aging effect of optic nerve injury and promoted optic nerve regeneration (Lu et al., 2020). Ezh2 is also critical for efficient cellular reprogramming (Ding et al., 2014; Onder et al., 2012), and has been shown to play a role in shaping the aging epigenome (Mozhui and Pandey, 2017). Moreover, we showed that Ezh2 overexpression specifically silenced the transcription of many genes functionally involved in synaptic activities of mature neurons, in some way rejuvenating RGCs at the transcriptional level. We thus wondered if *Ezh2* overexpression could also reverse the epigenetic aging effect of ONC on RGCs. We intravitreally injected AAV2-*GFP*, AAV2-*Ezh2*, or AAV2-*Ezh2-Y726D* into mice of exactly the same age, performed an ONC two weeks after the injection, and extracted DNA from FACS-enriched RGCs three days after the ONC. Control mice were only injected with AAV2 vectors but did not undergo the ONC. Reduced representation bisulfite sequencing libraries were constructed from RGC DNA and sequenced to obtain the DNA methylation landscape. We used a predictive PCA model described in (Lu *et al*., 2020) to estimate DNA methylation aging changes in RGCs. The results confirmed that three days after the ONC, epigenetic aging of RGCs was accelerated as indicated by DNA methylation aging signature (Fig. 8A). However, neither wild type Ezh2 nor Ezh2-Y726D could reverse the changes caused by the ONC (Fig. 8A), indicating that *Ezh2* overexpression in RGCs did not change the DNA methylation aging clock. In support of this, our RNA-seq results did not detect significant changes in mRNA levels of most 5mC DNA methyltransferases and demethylases (Fig. 8B, C). The different aging representations in the DNA methylation clock and transcriptomic landscape were not surprising. A recent study (Horvath et al., 2022) of naked mole rats (NMR), which live an exceptionally long life and are considered to be a nonaging mammal, showed normal aging progress of many tissues at the epigenetic level without significant overlap with age-related transcriptomic changes.

**Figure 8.**
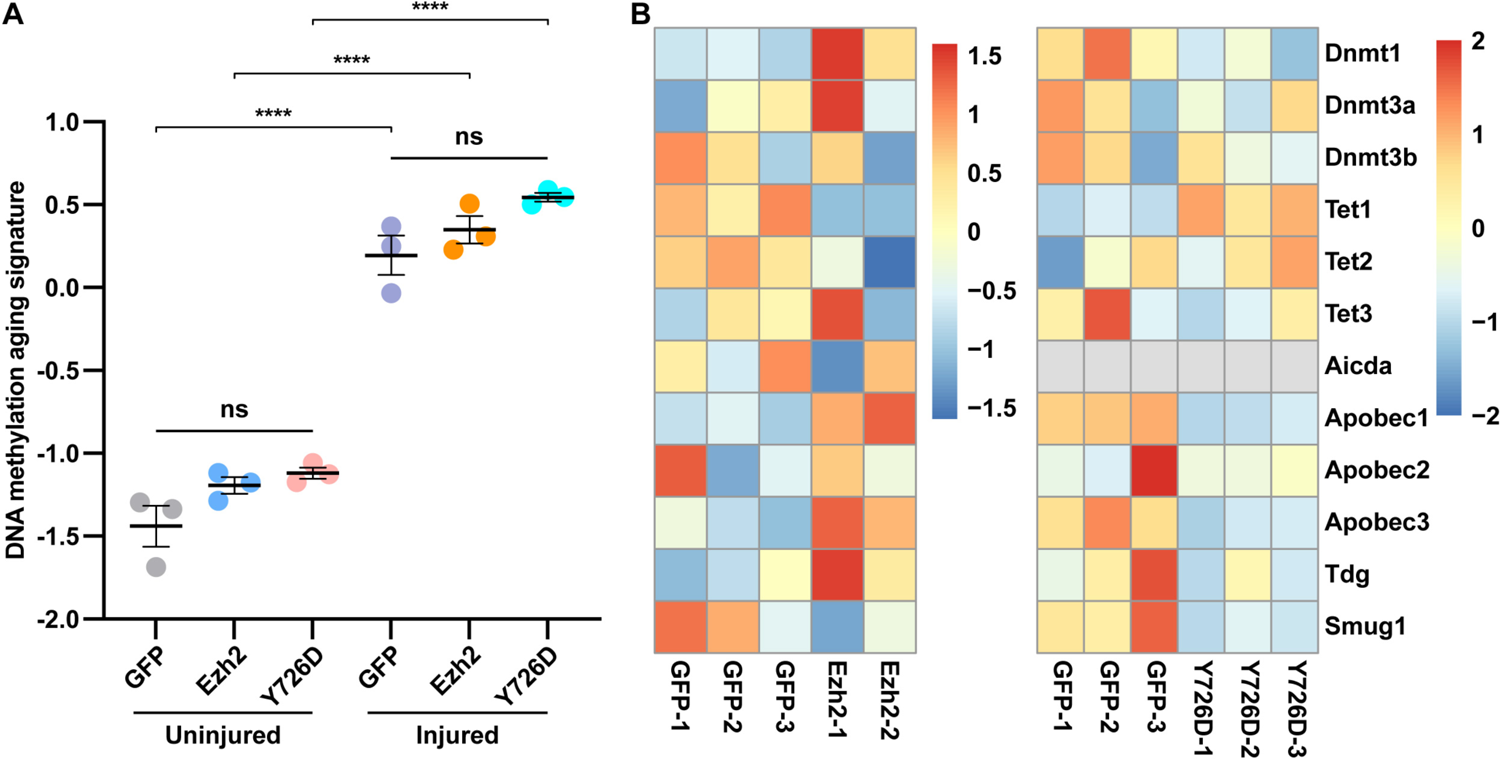
*Ezh2* overexpression does not alter the epigenetic aging clock of RGCs. (A) DNA methylation aging signature of RGCs is increased by optic nerve injury, but not affected by *Ezh2* or *Ezh2-Y726D* overexpression (one-way ANOVA followed by Tukey’s multiple comparisons; p < 0.0001; n = 3 reduced representation bisulfite sequencing libraries in each condition). (B) Heatmaps of mRNA levels of 5mC DNA methyltransferases and demethylases showing that most of them are not significantly changed by *Ezh2* or *Ezh2-Y726D* overexpression in RGCs. Note that *Aicda* mRNA was not detected in the control vs. *Ezh2-Y726D* overexpression RNA-seq. Data are represented as mean ± SEM. *P values* of post hoc analyses are illustrated in the figure. ****p < 0.0001.

Collectively, our study not only revealed novel roles of Ezh2 in coordinating axon regeneration via epigenetically regulation of multiple key regenerative pathways, but also suggested chromatin accessibility as a promising new target to promote CNS axon regeneration.

## Discussion

### Developmental decline of axon regeneration ability and transcriptomic rejuvenation

Axon regeneration ability of mammalian neurons declines as they mature. While PNS neurons can reactivate such an ability upon peripheral axonal injury, most adult CNS neurons permanently lose their ability to regenerate axons. Given that every single cell in an organism has exactly the same genome, and so does a neuron in different states (e.g., young vs. mature, healthy vs. injured), it is conceivable that the tuning of the axon regeneration ability in neurons is largely regulated by changes in the epigenomic and transcriptomic landscapes. Here we found that Ezh2, an epigenetic regulator that controls chromatin accessibility and gene transcription via histone methylation, was developmentally downregulated in both the PNS and CNS and could be upregulated in PNS neurons by peripheral nerve injury. Thus, Ezh2 level in the nervous system and the axon growth and regeneration potential of neurons rise and fall in parallel. Indeed, we found *Ezh2* loss-of-function impaired the normal axon regeneration of PNS neurons, while *Ezh2* gain-of-function promoted axon regeneration in non-regenerative adult CNS neurons. Mechanistic exploration with RNA-seq revealed that *Ezh2* overexpression in RGCs suppressed the transcription of a large number of genes regulating synaptic transmission and neuronal excitability, which are housekeeping functions of mature neurons. Therefore, our study suggested that Ezh2 upregulation might rejuvenate mature neurons at the transcriptomic level to empower them with stronger axon regeneration ability. In support of this, several previous studies also demonstrated that suppressed transcription of synaptic function-related genes positively correlated with enhanced axon regeneration ability (Hilton *et al*., 2022; Norsworthy et al., 2017; Sekine et al., 2018; Wang et al., 2018b). On the other hand, *Ezh2* overexpression also resulted in upregulation of many factors known to enhance axon regeneration, some of which are highly expressed in young developing neurons (Costales and Kolevzon, 2016; Penzo-Mendez, 2010; Zaytseva et al., 2020).

When the epigenetic aging biomarker, the DNA methylation clock, was examined in RGCs (Lu *et al*., 2020), we confirmed that optic nerve injury significantly accelerated RGC epigenetic aging. However, *Ezh2* overexpression was not able to reverse it. These results indicated that physiological aging at the transcriptomic level could be uncoupled from the DNA methylation aging clock. Indeed, a new study of NMR (Horvath *et al*., 2022), a species that lives an exceptionally long life with very slow age-related decline in physiological capacity, revealed by DNA methylation clocks that many types of tissue cells of these animals still showed aging effects at the epigenetic level. However, age-related DNA methylation changes did not overlap with age-related transcriptomic changes in liver cells of the NMR. Interestingly, iPS reprogramming was able to rejuvenate the DNA methylation clocks of NMR cells, consistent with a recent study in which partial reprogramming of RGCs with 3 reprogramming factors, Oct4, Sox2, and Klf4, reversed the DNA methylation aging induced by optic nerve injury (Lu *et al*., 2020). Collectively, we think that *Ezh2* overexpression in mature RGCs switched their transcriptomic landscape back to a younger stage with stronger axon growth ability. The ineffectiveness of Ezh2 on DNA methylation clock was likely due to the unaltered levels of most DNA methylation-related enzymes (see Fig. 8).

### Ezh2 is a master suppressor of CNS axon regeneration inhibitory signaling

It is well recognized that there are three major classes of axon regeneration inhibitors in the extracellular environment of the mature CNS, including MAIs, CSPGs, and repulsive axon guidance cues (Giger *et al*., 2010). Here we showed that that *Ezh2* overexpression silenced transcription of OMgp, which is one of the three major MAIs, Lingo3, which is a potential co-receptor of MAIs and CSPGs, and several ephrin receptors, which are chemorepellent axon guidance cues during both development and regeneration. Specifically, OMgp is one of the three major MAIs (OMgp, MAG, Nogo) that act through Nogo receptor 1 (NgR1) complex or PirB to inhibit axon growth (Atwal et al., 2008; Fournier et al., 2001; Wang et al., 2002). Despite its name, OMgp is also expressed by neurons (Habib et al., 1998). Although *Lingo3* has not been directly linked to mammalian axon regeneration, its paralog *Lingo1* encodes a critical component of the NgR1 complex (Mi et al., 2004), which functions to block axon regeneration via RhoA pathway when activated by myelin-associated inhibitors (MAIs) and CSPGs (Dickendesher et al., 2012). *Lingo1* loss-of-function promotes axon regeneration and neuronal survival in various CNS injury and disease models (Fu et al., 2008; Inoue et al., 2007; Ji et al., 2006). A recent study showed that Lingo family receptors could form heteromers with one another in the mouse brain (Guillemain et al., 2020), suggesting potential functional overlap between the paralogs. Tenascin-R is an extracellular matrix molecule that was shown in previous studies to be a repulsive guidance cue in zebrafish during development (Becker et al., 2003) and an inhibitor of mouse optic nerve regeneration (Becker et al., 2000). Ephrin receptors are chemorepellent axon guidance molecules that can cause growth cone collapse when activated by their ligands, ephrins (Taylor et al., 2017). Among them, EphA4 has been shown to be upregulated following spinal cord injury and act to inhibit axon regeneration in rodents (Giger *et al*., 2010). Functionally, we provided direct and strong evidence that overexpression of *Omg* or *Lingo3* blocked optic nerve regeneration induced by *Ezh2* overexpression to a great extent. Thus, apart from tuning down synaptic function-related gene transcription in mature neurons, Ezh2 also acted to transcriptionally repress the expression of either CNS axon regeneration inhibitors or their receptors, suggesting that it might be a master suppressor of CNS axon regeneration inhibitory signaling pathways. Notably, our study indicated that besides glial cells, neurons themselves could also contribute to the production of extracellular CNS regeneration inhibitors, such as OMgp and tenascin-R. It would be interesting for future studies to investigate the autocrinal mechanisms of neuron-secreted axon regeneration inhibitors.

### Ezh2 enhances optic nerve regeneration via both methylation-dependent and -independent pathways

As a protein with a methyltransferase domain, Ezh2 was originally identified as the catalytic core protein of the histone modification complex PRC2. Thus, most previous studies have focused on investigating the roles of Ezh2 or PRC2-mediated H3K27 trimethylation and transcriptional repression in various biological events. Evidence has been, however, emerging to suggest that Ezh2 also has protein methylation-unrelated activities. For example, a prostate cancer study showed that Ezh2 could transactivate androgen receptor by directly binding to its promoter region (Kim *et al*., 2018). Likewise, a breast cancer study found that a ternary complex of Ezh2, RelA, and RelB bound to promoters of *Il6* and *Tnf* to enhance their transcription (Lee *et al*., 2011). Furthermore, a more recent study discovered that Ezh2 was even able to regulate protein translation via interacting with fibrillarin and controlling rRNA methylation in a way independent of its methyltransferase activity (Yi et al., 2021). In our study, an Ezh2 mutant lacking the methyltransferase activity (Ezh2-Y726D) was still able to weakly promote optic nerve regeneration, strongly indicating that methyltransferase-independent activities of Ezh2 also contributed to the enhanced axon regeneration. In the current study, we did not further investigate these methyltransferase-independent mechanisms.

Our RNA-seq results provided some evidence that the catalytically dead Ezh2-Y726D acted in a dominant negative manner to control the general patterns of gene transcription compared to wild type Ezh2. Such results seemed confusing because Ezh2-Y726D did promote RGC survival and optic nerve regeneration. One likely explanation is that under the control condition, optic nerve regeneration does not occur due to the lack of intrinsic axon growth ability of mature RGCs. The overall effect of Ezh2-Y726D expression is to enhance the maturation state of RGCs with increased expression of synaptic function-related genes, which is unlikely to further reduce the already very low intrinsic axon growth ability. However, our results did show that Ezh2-Y726D regulated some genes in the same way as did Ezh2 (see Fig. 4F, H), which might provide some clues of how this Ezh2 mutant promoted optic nerve regeneration. For example, both *Ezh2* and *Ezh2-Y726D* overexpression significantly upregulated the mRNA level of neuropeptide Y (*Npy*), which is a traditional regeneration-associated gene (Ma and Willis, 2015) and has been reported to attract growth cones and promote axon growth of embryonic rat DRG neurons (Sanford et al., 2008). In addition, a recent study discovered that *Npy* knockdown impaired supraspinal axon regeneration and locomotor function recovery in adult zebrafish after spinal cord injury (Cui et al., 2021). Likewise, both Ezh2 and Ezh2-Y276D downregulated mRNA levels of corticotropin releasing hormone binding protein (*Crhbp*) (Table S1) and *Slc6a13* (see Fig. 4F and Table S1). *Crhbp* was shown to be selectively expressed in RGC subtypes susceptible to ONC (Tran *et al*., 2019). Loss-of-function of *Crhbp* significantly promoted RGC survival and optic nerve regeneration (Tran *et al*., 2019). Our current study showed that *Slc6a13* negatively regulated axon regeneration in both the PNS and CNS (see Fig. 5B, C and Fig. S6A, B). Upregulation of *Npy* and/or downregulation of *Crhbp* and *Slc6a13* might be methylation-independent mechanisms by which Ezh2-Y726D modestly induced optic nerve regeneration. Future studies are needed to further explore non-canonical roles of Ezh2 in mammalian axon regeneration.

Although AAV2 vectors also transduce other retinal cells besides RGCs, our data showed that Ezh2-induced upregulation of H3K27me3 was mostly observed in cells within the ganglion cell layer of the retina (see Fig. 2C), which contains mainly RGCs and some displaced amacrine cells. However, we cannot exclude the possibility that the optic nerve regeneration induced by *Ezh2* overexpression was also contributed by non-RGC-autonomous factors (Zhang et al., 2019). Future studies using the Vglut2-Cre mouse line to specifically restrict *Ezh2* expression in RGCs will provide a clear answer.

## Acknowledgments

We acknowledge Dr. Michele Pucak from the Multiphoton Imaging Core (supported by NS050274) of the Department of Neuroscience, Johns Hopkins School of Medicine and Hao Zhang from the Flow Cytometry Cell Sorting Core Facility at Bloomberg School of Public Health, Johns Hopkins University for their help in confocal microscopy and FACS sorting. We appreciate the Johns Hopkins Single Cell & Transcriptomics Core for RNA-seq and ATAC-seq experiments. F.-Q.Z. has been supported by grants from NIH (R01NS085176, R01EY027347, R01EY030883, R01EY031779), the Craig H. Neilsen Foundation (259450), and the BrightFocus Foundation (G2017037). X.-W.W is supported by an NIH grant (K99EY031742). J.Q. is supported by NIH (R01EY029548).

## Author contributions

X.-W.W., C.-M.L., and F.-Q.Z. conceived the study and designed the project; X.-W.W., S.-G.Y., and R.-Y-W. performed the experiments; M.-W.H., X.L., and J.Q. analyzed the sequencing data; X.-W.W. and C.Z. performed data analysis; J.-J.J, A.R.K., and A.J.O helped with the data analysis; X.-W.W. and F.-Q.Z. wrote the manuscript with contributions from all authors.

## STAR Methods

### RESOURCE AVAILABILITY

#### Lead contact

Further information and requests for resources and reagents should be directed to and will be fulfilled by the Lead Contact, Feng-Quan Zhou.

#### Materials availability

All mouse lines and plasmids generated in this study are available from the lead contact with a completed materials transfer agreement.

#### Data and code availability

The data that support the findings of this study are available from the corresponding author upon reasonable request. All sequencing raw data and processed data will be deposited to Gene Expression Omnibus upon publication.

## EXPERIMENTAL MODEL AND SUBJECT DETAILS

### Adult DRG neuronal culture

Lumbar DRGs were dissected from euthanized 6-8-week-old mice, digested with 1 mg/ml type I collagenase (Thermo Fisher Scientific) and 5 mg/ml dispase II (Thermo Fisher Scientific) at 37°C for 70 min, washed 3 times with HBSS, and dissociated into cell suspension by repeated pipetting in MEM containing 10% fetal bovine serum (FBS) and 1% penicillin/streptomycin (pen/strep, Thermo Fisher Scientific). Cells were filtered with a 100-μm cell strainer and pelleted by centrifugation at 500 rcf for 5 min.

For electroporation experiments, pelleted cells were resuspended with 100 μl nucleofection buffer (Mouse Neuron Nucleofector Kit, Lonza) containing siRNA oligos (0.2 nmol) and/or plasmid vectors (10 μg) and electroporated with Nucleofector II (Lonza). Cells were then immediately resuspended in pre-warmed (37°C) MEM containing 5% FBS, 1% GlutaMAX-I (Thermo Fisher Scientific), 1% pen/strep and antimitotic reagents (20 μM 5-Fluoro-2’-deoxyuridine and 20 μM uridine, both from Sigma-Aldrich), plated on coverslips pre-coated with 100 μg/ml poly-D-lysine (Sigma-Aldrich) and 10 μg/ml laminin (Thermo Fisher Scientific), and cultured for 3 days at 37°C in a humidified incubator containing 5% CO_2_. Culture medium was refreshed 6 hours after plating.

For replate experiments, electroporated cells were plated on pre-coated dishes and cultured for 3 days. Cells were then forced to detach from dishes by pipetting, replated on pre-coated coverslips, and cultured for 24 hours.

For culture of conditioning-lesioned DRG neurons, lumbar 4 and 5 DRGs were dissected from *Ezh2^f/f^* and *Advillin-Cre; Ezh2^f/f^* mice 3 days after sciatic nerve transection (see **Sciatic nerve crush or transection**). After enzymatic digestion and dissociation, filtered cells were immediately plated on pre-coated coverslips and cultured for 24 hours.

### Mice

All animal experiments were conducted in accordance with the protocol approved by the Institutional Animal Care and Use Committee of the Johns Hopkins University. Adult C57BL/6J mice (6-10 weeks) of both sexes were used unless otherwise stated. The *Ezh2^f/f^* (stock# 015499-UNC) mouse strain was obtained from the Mutant Mouse Resource and Research Center (MMRRC) at University of North Carolina at Chapel Hill, an NIH-funded strain repository, and was donated to the MMRRC by Alexander Tarakhovsky, Ph.D., The Rockefeller University. The *Advillin-Cre* mouse line (JAX stock# 032536) was a kind gift from Dr. Fan Wang’s laboratory at Duke University and was crossed with *Ezh2^f/f^* to obtain *Advillin-Cre; Ezh2^f/f^* conditional knockout mice. Genotypes of the mice were determined by PCR using programs and primers provided by the MMRRC and Jackson Laboratory. All mouse surgeries were performed under anesthesia induced by intraperitoneal injection of ketamine (100 mg/kg) and xylazine (10 mg/kg) diluted in sterile saline. Details of the surgeries are described below.

## METHOD DETAILS

### Constructs

The mouse *Ezh2* open reading frame (ORF) was cloned into *pAAV-CMV* with a 5’ NheI restriction site and a 3’ XhoI restriction site to obtain *pAAV-CMV-Ezh2*. *pAAV-CMV-Ezh2-Y726D* was constructed by mutating the 2,176^th^ nucleotide of *Ezh2* ORF from a T to a G. The mouse *Slc6a13* ORF with a 5’ NheI restriction site and a 3’ NotI restriction site was cloned into *pAAV-CMV* to obtain *pAAV-CMV-Slc6a13* or used to replace the *EGFP* ORF in *pCMV-EGFP* to obtain *pCMV-Slc6a13*. Mouse *Lingo3*, *Omg*, and *Wfdc1* ORFs were cloned into *pAAV-CMV* using a 5’ KpnI restriction site and a 3’ ApaI restriction site to obtain *pAAV-CMV-Lingo3*, *pAAV-CMV-Omg*, and *pAAV-CMV-Wfdc1*, respectively. The mouse *Neurog2* ORF flanked by a 5’ BamHI restriction site and a 3’ EcoRV restriction site was used to replace the *EYFP* open reading frame in *pAAV-Ef1a-EYFP* to obtain *pAAV-Ef1a-Neurog2*. Mouse *Lingo3* and *Neurog2* ORFs were synthesized by Integrated DNA Technologies with codon optimization. All restriction enzymes and the T4 DNA ligase were purchased from New England Biolabs. Plasmids were amplified using DH5α competent cells (Thermo Fisher Scientific) and purified with the Endofree Plasmid Maxi Kit (Qiagen). AAV2-*GFP* (SL100812) was purchased from SignaGen Laboratories. All other AAV2 viruses were also packaged by SignaGen Laboratories.

### Immunocytochemistry

Cultured DRG cells were fixed with 4% PFA for 15 min at room temperature, washed 3 times with PBS, and blocked with PBST (0.3%) containing 10% goat serum for 1 hour at room temperature. After blocking, cells were incubated in anti-tubulin β3 (1:1,000, BioLegend) and/or anti-GFP (1:1,000, Thermo Fisher Scientific) primary antibodies for 1 hour at room temperature, washed 3×10 min with PBS, incubated in corresponding Alexa Fluor-conjugated secondary antibodies (1:500, Thermo Fisher Scientific) for 1 hour at room temperature, and washed 3×10 min again with PBS. All antibodies were diluted with PBST (0.3%) containing 10% goat serum. Coverslips were then mounted in Fluoroshield (Sigma-Aldrich) onto microscope slides.

### Analysis of *in vitro* DRG neuron axon growth

Fluorescent images of cultured DRG neurons were obtained with a Zeiss inverted fluorescence microscope controlled by the AxioVision software using a 5× objective. The longest axon of each neuron was manually traced and measured with the built-in “measure/curve spline” function of the AxioVision software. Only neurons with axons longer than twice the diameter of their somas were included. In most experiments, at least 60 neurons were analyzed in each condition. Measurement was done by experimenters blinded to experimental conditions.

### *in vivo* DRG electroporation

Under anesthesia, a small dorsolateral laminectomy was performed on the left side to expose left lumbar 4 and 5 DRGs. Using a pulled glass micropipette (World Precision Instruments) connected to a Picospritzer III (pressure: 20 psi, pulse duration: 6 ms, Parker Hannifin), 1 μl plasmid vectors (2 μg) and/or siRNA oligos (0.1 nmol) containing 0.05% fast green FCF (Sigma-Aldrich) were injected into each DRG. After injection, *in vivo* electroporation was performed by applying five electric pulses (voltage: 35 V, pulse duration: 15 ms, pulse interval: 950 ms) using a platinum tweezertrode (BTX) powered by an ECM 830 Electro Square Porator (BTX). The wound was then closed with sutures.

### Sciatic nerve crush or transection

Under anesthesia, sciatic nerves were exposed right below the pelvis and crushed with Dumont #5 forceps (Fine Science Tools) for 15 s or cut with scissors, and the wound was closed by sutures. In sham surgeries, sciatic nerves were only exposed but not injured. Sciatic nerve crush was used in *in vivo* DRG neuron axon regeneration experiments and was only performed on the left side. The crush site was marked with 10-0 nylon epineural sutures. Sciatic nerve transection was used in other experiments and was performed bilaterally.

### Analysis of *in vivo* DRG neuron axon regeneration

Three or five days after sciatic nerve crush, mice were anesthetized and transcardially perfused with PBS followed by 4% PFA. Sciatic nerve segments (proximal end: 5 mm proximal to the crush site, distal end: the point where the sciatic nerve branches into three nerves) were dissected and post-fixed in 4% PFA overnight at 4°C. On the next day, nerve segments were mounted in Fluoroshield (Sigma-Aldrich) onto microscope slides, covered with coverslips, and flattened by applying a heavy weight on coverslips. Tiled fluorescent images of whole-mount nerve segments were obtained with a Zeiss inverted fluorescence microscope controlled by the AxioVision software using a 5× objective. Using the built-in “measure/curve spline” function of the AxioVision software, GFP-labeled axons were manually traced from the crush site to axonal tips to determine the lengths. The mean length of all axons traced in one nerve segment was used as the average axon length of this nerve. Nerves whose epineural sutures were missing or with less than 10 identifiable GFP-labeled axons were excluded from data analysis. Measurement was done by experimenters blinded to experimental conditions.

### Optic nerve crush and regeneration model

Intravitreal virus injection, optic nerve crush and RGC axon labeling were performed as previously described (Wang *et al*., 2020). Briefly, under anesthesia, 1.5 μl AAV2 virus (∼1×10^13^ genome copies/ml) was injected into the vitreous humor with a Hamilton syringe (33-gauge needle). The position and direction of the needle were well controlled to avoid injury to the lens. Two weeks after the virus injection, under anesthesia, a small incision was made in the skin right behind the eye and the conjunctiva was incised to expose the extraocular muscles. The muscles were pushed aside with forceps to expose the optic nerve, and the optic nerve was crushed with Dumont #5 forceps (Fine Science Tools) for 5 s at approximately 0.5 mm behind the optic disc. Care was taken to avoid damage to the ophthalmic artery. For the post-injury treatment model, the optic nerve crush was done 1 day prior to the virus injection. To label RGC axons in the optic nerve, under anesthesia, 1.5 μl Alexa Fluor 555 or 647-conjugated CTB (1 μg/μl, Thermo Fisher Scientific) was injected into the vitreous humor with a Hamilton syringe (33-gauge needle) 2 days before tissue harvesting.

### Optic nerve tissue clearing

Two days after intravitreal CTB injection, mice were anesthetized and transcardially perfused with PBS followed by 4% PFA. Optic nerves were dissected and post-fixed in 4% PFA overnight at 4°C. On the next day, optic nerves were washed 3 times with PBS, dehydrated in an ascending series of tetrahydrofuran (50%, 70%, 80%, 100% and 100%, v/v in distilled water, 20 min each, Sigma-Aldrich), and cleared in a 1:2 mixed solution of benzyl alcohol and benzyl benzoate (BABB, Sigma-Aldrich). Incubations were done on an orbital shaker at room temperature. Nerves were stored in BABB and protected from light at room temperature before imaging.

### Analysis of optic nerve regeneration

Z-stacked (step size: 2 μm) and tiled fluorescent images of tissue-cleared whole-mount optic nerves were obtained with a Zeiss LSM 800 confocal microscope using a 20× objective. To quantify the number of regenerating axons in each optic nerve, every 8 consecutive planes were Z-projected in maximum intensity to generate a series of Z-projection images of 16-μm-thick optical sections. At each 250-μm interval from the crush site, the number of CTB-labeled axons was counted in each Z-projection image and summed over all optical sections.

### NMDA-induced excitotoxicity model

Under anesthesia, 1.5 μl AAV2 virus (∼1×10^13^ genome copies/ml) was injected into the vitreous humor with a Hamilton syringe (33-gauge needle). Two weeks after the virus injection, under anesthesia, 1.5 μl NMDA (20 mM, Sigma-Aldrich) was injected into the vitreous humor with a Hamilton syringe (33-gauge needle). The position and direction of the needle were well controlled to avoid injury to the lens.

### RGC enrichment by fluorescence-activated cell sorting (FACS)

Retinas were dissected from euthanized mice, digested with papain (Thermo Fisher Scientific) containing 0.005% DNase (Worthington) at 37°C for 8 min, washed 3 times with HBSS, and dissociated into cell suspension by repeated pipetting in NeuroBasal medium containing 1% BSA. Cells were filtered with a 40-μm cell strainer, pelleted by centrifugation at 500 rcf for 5 min, resuspended in NeuroBasal medium containing 1% BSA, blocked with mouse CD16/CD32 Fc block antibody (1:50) for 5 min on ice, and labeled with PE-conjugated anti-CD90.2 antibody (1:100) for 30 min on ice. After that, cells were washed twice with HBSS containing 1% BSA, pelleted by centrifugation at 500 rcf for 5 min, and again resuspended in Neurobasal medium containing 1% BSA. Propidium iodide (PI) or DAPI was mixed with the cell suspension to label dead cells 2 min before cells were loaded into a Beckman Coulter MoFlo Legacy Cell Sorter. CD90.2-positive and PI or DAPI-negative cells were identified and sorted into NeuroBasal medium containing 1% BSA with a 70-μm nozzle.

### RNA sequencing and data analysis

RNA was isolated from FACS-enriched RGCs 3 days after optic nerve crush using the PicoPure RNA Isolation Kit (Thermo Fisher Scientific) following the manufacture’s manual. RNA quality was verified using the Agilent Fragment Analyzer (Agilent Technologies). RNA sequencing libraries were prepared using the TruSeq Stranded Total RNA Kit (Illumina) and quality checked by the Agilent Fragment Analyzer. Equimolar amounts of the finished libraries were then pooled and sequenced on an Illumina NextSeq 500 with 2×75 bp paired reads.

Raw FASTQ data were mapped to mouse reference genome (GRCm38) using STAR aligner (version 2.7.0d) with default parameters. The number of counts per gene was estimated using the “quantMode” command in STAR. Quantified raw counts were used in DESeq2 (version 1.22.2) for the analysis of differentially expressed genes (DEGs). Genes with less than 10 counts in total from six libraries were excluded from analysis. Genes with adjusted *P* < 0.05 and fold change > 1.5 were chosen as DEGs. Principle component analysis and clustering analysis were also performed with the transformed count matrix in DESeq2. Normalized counts were used to produce heatmaps (Fig. 4F-H, Fig. 8B, Fig. S3E) and scatter plots (Fig. S4D-G). GO analysis (biological process) was done using DAVID Bioinformatics Resources 6.8.

### Assay for transposase-accessible chromatin with sequencing (ATAC-seq) and data analysis

ATAC-seq libraries were constructed from FACS-enriched RGCs (50,000 cells for each library) 3 days after the optic nerve crush following a previously published protocol (ref). Briefly, cells were pelleted by centrifugation at 500 rcf for 5 min, washed with ice-cold PBS, pelleted again by centrifugation at 500 rcf for 5 min, and lysed in 50 μl ice-cold lysis buffer. Immediately after lysis, nuclei were pelleted by centrifugation at 500 rcf for 10 min, resuspended in 50 μl transposase reaction mix and incubated at 37°C for 30 min. After the transposition reaction, the product was purified with the DNA Clean & Concentrator-5 Kit (Zymo Research). 20 μl tagmented DNA was PCR amplified with NEBNext High-Fidelity PCR Master mix and forward and reverse UDI primers. Amplification was first performed for 5 cycles, following which 5 μl of each partially amplified library was used to perform qPCR to determine the additional number of PCR cycles needed for each library. Final amplified libraries were purified using 1.1× Ampure XP bead purification. Equimolar amounts of the finished libraries were then pooled and sequenced on an Illumina NovaSeq 6000 with 2×100 bp paired reads.

After removing adapters of Illumina reads with Cutadapt, pair-end ATAC-Seq reads were mapped to the mouse reference genome (GRCm38) using Bowtie2 with default parameters. Qualified properly paired reads (MAPQ score > 10) were assessed by SAMTools. Duplicate reads were removed with MarkDuplicates function from Picard. After using MACS2 to call peak regions of each sample, we used multiBamSummary function in deepTools to calculate read counts for all samples. ChIPseeker was used to annotate genomic context of identified peaks. Gene annotation information was accessed using TxDb.Mmusculus.UCSC.mm10.knownGene. Differential accessibility analysis was performed by DESeq2.

### Cleavage under targets and tagmentation (CUT&Tag) and qPCR

ChIP-seq libraries were constructed from FACS-enriched RGCs (100,000 cells for each library) using an H3K27me3 antibody (1:100, Active Motif) or normal rabbit IgG (1:100, Sigma-Aldrich) and the CUT&Tag-IT Assay Kit (Active Motif) following the manufacture’s manual. Identical amount of DNA from each library was used in qPCR to determine the enrichment of locus sequences in the promoter region of each gene. Fold enrichment of DNA bound to H3K27me3 was determined using the ddCt method and normalized to IgG. All qPCR experiments were done in triplicate.

### Reduced representative bisulfite sequencing (RRBS) and data analysis

Three days after the optic nerve crush (injured conditions) or 17 days after AAV2 injection (uninjured conditions), DNA was extracted from FACS-enriched RGCs using the Quick-DNA Microprep Plus Kit (Zymo Research) following the manufacture’s manual. 10 ng genomic DNA was digested with 30 units of MspI (NEB). Fragments were ligated to pre-annealed adapters containing 5’-methyl-cytosine instead of cytosine according to Illumina’s specified guidelines. Adaptor-ligated fragments ≥ 50 bp in size were recovered using the DNA Clean & Concentrator-5 Kit (Zymo Research). The fragments were then bisulfite-treated using the EZ DNA Methylation-Lightning Kit (Zymo Research). Preparative-scale PCR was performed, and the product was purified with the DNA Clean & Concentrator-5 Kit (Zymo Research) for sequencing on an Illumina NovaSeq with 2×150 bp paired reads.

Sequencing reads from bisulfite-treated classic RRBS libraries were identified using standard Illumina base calling software and then raw FASTQ files were adapter, filled-in nucleotides, and quality trimmed using TrimGalore 0.6.4. FastQC 0.11.8 was used to assess the effect of trimming and overall quality distributions of the data. Alignment to the mm10 reference genome was performed using Bismark 0.19.0. Methylated and unmethylated read totals for each CpG site were called using MethylDackel 0.5.0. The methylation level of each sampled cytosine was estimated as the number of reads reporting a C, divided by the total number of reads reporting a C or T.

DNA methylation aging signature was estimated by the method described in (Lu *et al*., 2020). We used 1-month, 12-month, and 18-month mouse samples in that study as the training set in the predictive PCA model. Differential CpG sites were selected based on top 50% CpGs using biweight midcorrelation among three comparisons (i.e., AAV2-*GFP* uninjured vs AAV2-*GFP* injured, AAV2-*Ezh2* uninjured vs AAV2-*Ezh2* injured, and AAV2-*Ezh2-Y726D* uninjured vs AAV2-*Ezh2-Y726D* injured). The first principal component was chosen for representing the DNA methylation aging signature.

### Immunoblotting

Total protein was extracted from mouse DRGs, cultured DRG cells, retinas, or FACS-enriched RGCs using the RIPA buffer (Sigma-Aldrich) containing the protease inhibitor cocktail (Sigma-Aldrich) and phosphatase inhibitor cocktail (Sigma-Aldrich). Identical amount of total protein from each condition was separated by SDS-PAGE on 4-12% Bis-Tris gels and transferred onto polyvinylidene difluoride membranes. Membranes were blocked with TBST containing 5% blotting-grade blocker (Bio-Rad), incubated in primary antibodies against target molecules overnight at 4 °C, washed 4 times (5, 5, 10, 10 min) with TBST, incubated in corresponding HRP-linked secondary antibodies (1:2,000, Cell Signaling Technology) for 1 hour at room temperature, and washed 4 times (5, 5, 10, 10 min) again with TBST. All antibodies were diluted with TBST containing 5% blotting-grade blocker. Rabbit anti-Ezh2 (1:1,000) and anti-H3 (1:1,000) were purchased from Cell Signaling Technology. Mouse anti-H3K27me3 (1:10,000), anti-β-actin (1:10,000), and anti-Gapdh (1:20,000) were purchased from Sigma-Aldrich.

### Immunohistochemistry of whole-mount retinas

Retinas were dissected from transcardially perfused mice and post-fixed in 4% PFA overnight at 4°C. On the next day, retinas were post-fixed in ice-cold methanol for 20 min, washed 3 times with PBS, radially cut into a petal shape, and blocked with PBST (1%) containing 10% goat serum for 1 hour at room temperature. After blocking, retinas were incubated in primary antibodies against target molecules overnight at 4°C, washed 4×15 min with PBST (0.3%), incubated in corresponding Alexa Fluor-conjugated secondary antibodies (1:500, Thermo Fisher Scientific) for 2 hours at room temperature, and washed 4×15 min again with PBST (0.3%). All antibodies were diluted with PBST (1%) containing 10% goat serum. Retinas were flat-mounted in Fluoroshield (Sigma-Aldrich) onto microscope slides and covered by coverslips. Fluorescent images of whole-mount retinas were obtained with a Zeiss LSM 800 confocal microscope using a 20× objective.

### Analysis of RGC survival rate

To quantify RGC survival rate, mice were transcardially perfused 2 weeks after optic nerve crush or 1 week after NMDA injection and both retinas of each mouse were dissected. Retinas were stained with guinea pig anti-Rbpms (1:100, Thermo Fisher Scientific) following the steps described above (see **Immnunohistochemistry of whole-mount retinas**). 6-9 fields were randomly taken from the peripheral regions of each retina. For each mouse, RGC survival rate was calculated by dividing the average number of Rbpms-positive cells in one field in the injured retina by that in the uninjured retina. Only cells in the ganglion cell layer were counted.

### Immunohistochemistry of retinal sections

Fixed retinas were immersed in 30% sucrose overnight at 4°C. On the next day, retinas were embedded in the OCT compound, frozen, and cut into 10-μm sections with a cryostat. Sections were transferred onto microscope slides and warmed on a slide warmer for 1 hour at 37°C. Sections on slides were rinsed once in PBS, soaked in 100°C citrate buffer (pH 6) for 15 min, cooled down in the buffer to room temperature, washed 2×5 min with PBS, and blocked with PBST (0.3%) containing 10% goat serum for 1 hour at room temperature. After blocking, sections were incubated in primary antibodies against target molecules overnight at 4 °C, washed 4 times (5, 5, 10, 10 min) with PBST (0.3%), incubated in corresponding Alexa Fluor-conjugated secondary antibodies (1:500, Thermo Fisher Scientific) for 1 hour at room temperature, and washed 4 times (5, 5, 10, 10 min) again with PBST (0.3%). All antibodies were diluted with PBST (0.3%) containing 10% goat serum. Sections were mounted in Fluoroshield (Sigma-Aldrich) and covered by coverslips. Fluorescent images of retinal sections were obtained with a Zeiss inverted fluorescence microscope controlled by the AxioVision software using a 20× objective.

### Analysis of H3K27me3 levels in RGCs

To analyze H3K27me3 levels in RGCs, retinas were dissected from transcardially perfused mice 2 weeks after intravitreal injection of AAV2-*GFP*, AAV2-*Ezh2*, or AAV2-*Ezh2-Y726D* and sectioned. Retinal sections were stained with guinea pig anti-Rbpms (1:500, Thermo Fisher Scientific) and mouse anti-H3K27me3 (1:100, Sigma-Aldrich) following the steps described above (see **Immunohistochemistry of retinal sections**).

To quantify H3K27me3 levels in RGCs, fluorescence intensity of H3K27me3 immunoreactivity of at least 150 Rbpms-positive cells from 10-12 non-adjacent retinal sections acquired with identical imaging configurations was analyzed for each retina. Fluorescence intensity was measured using the “outline spline” function of the AxioVision software and the background fluorescence intensity was subtracted.

## QUANTIFICATION AND STATISTICAL ANALYSIS

Statistical analysis was done with GraphPad Prism 9 and the significance level was set as *P* < 0.05. Data are represented as mean ± SEM unless specifically stated. For comparisons between two groups, two-tailed unpaired or paired t test was used. For comparisons among three or more groups, one-way ANOVA followed by Tukey’s multiple comparisons was used to determine the statistical significance. All details regarding statistical analysis, including the tests used, *P* values, exact values of n, definitions of n, etc., are described in figure legends.

