## Supplemental figures for "Histone methyltransferase Ezh2 coordinates mammalian axon regeneration via epigenetic regulation of key regenerative pathways"

Figure S1

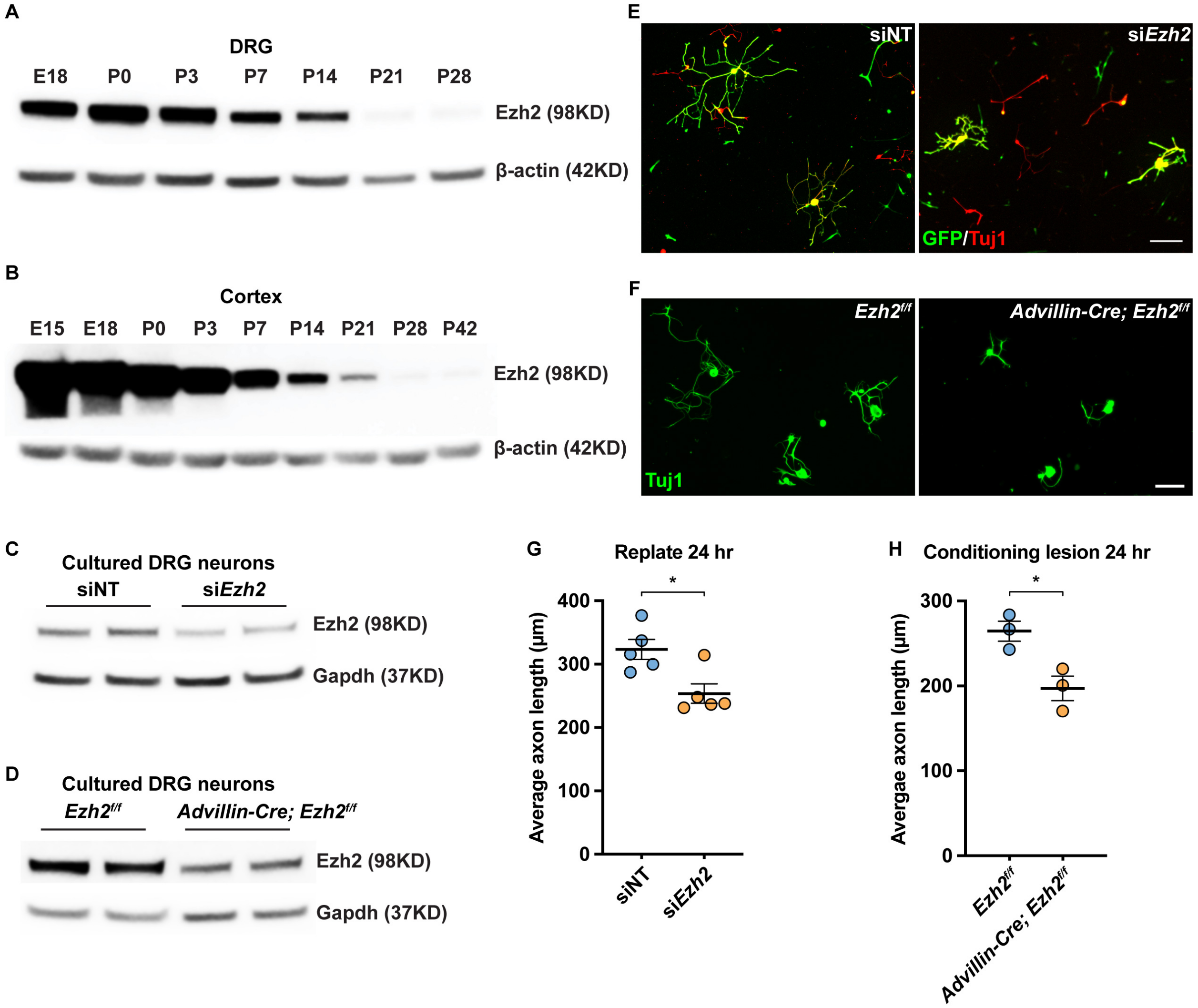

A
DRG
E18
P0
P3
P7
P14
P21
P28
Ezh2 (98KD)
β-actin (42KD)
B
Cortex
E15
E18
P0
P3
P7
P14
P21
P28
P42
Ezh2 (98KD)
β-actin (42KD)
C
Cultured DRG neurons
siNT
siEzh2
Ezh2 (98KD)
Gapdh (37KD)
D
Cultured DRG neurons
Ezh2f/f
Advillin-Cre; Ezh2f/f
Ezh2 (98KD)
Gapdh (37KD)
E
siNT
siEzh2
GFP/Tuj1
F
Ezh2f/f
Advillin-Cre; Ezh2f/f
Tuj1
G
Replate 24 hr
*
400
300
200
100
0
Average axon length (μm)
siNT
siEzh2
H
Conditioning lesion 24 hr
*
300
200
100
0
Avergae axon length (μm)
Ezh2f/f
Advillin-Cre; Ezh2f/f

**Figure S1. Ezh2 is developmentally downregulated in the nervous system and is necessary for axon growth of sensory neurons *in vitro.* Related to Figure 1.**

(A, B) Representative immunoblotting showing that Ezh2 is developmentally downregulated in the DRG (A, n = 2 independent experiments) and cortex (B, n = 3 independent experiments).

(C) Immunoblotting showing decreased Ezh2 level in cultured DRG neurons three days after electroporation of siRNAs targeting *Ezh2* mRNA (n = 2 in each condition).

(D) Immunoblotting showing decreased Ezh2 level in cultured DRG neurons of *Advillin-Cre; Ezh2*$^{f/f}$ mice (n = 2 in each condition).

(E) Representative immunocytochemistry of cultured DRG neurons showing that *Ezh2* knockdown impairs axon growth of sensory neurons *in vitro*. Cells were stained with anti-GFP (green) and anti-tubulin β3 (red). Scale bar, 200 µm.

(F) Representative immunocytochemistry of cultured DRG neurons showing that *Ezh2* knockout impairs axon growth of conditioning-lesioned sensory neurons *in vitro*. Cells were stained with anti-tubulin β3 (green). Scale bar, 100 µm.

(G) Quantification of the average length of the longest axon of each neuron in (E) (unpaired t test; p = 0.0132; n = 5 independent experiments; at least 60 neurons were analyzed for each condition in each independent experiment, except in one experiment only 26 neurons were analyzed in the *Ezh2* knockdown condition).

(H) Quantification of the average length of the longest axon of each neuron in (F) (unpaired t test; p = 0.0227; n = 3 independent experiments; at least 100 neurons were analyzed for each condition in each independent experiment).

Data are represented as mean ± SEM. *p < 0.05.

Figure S2

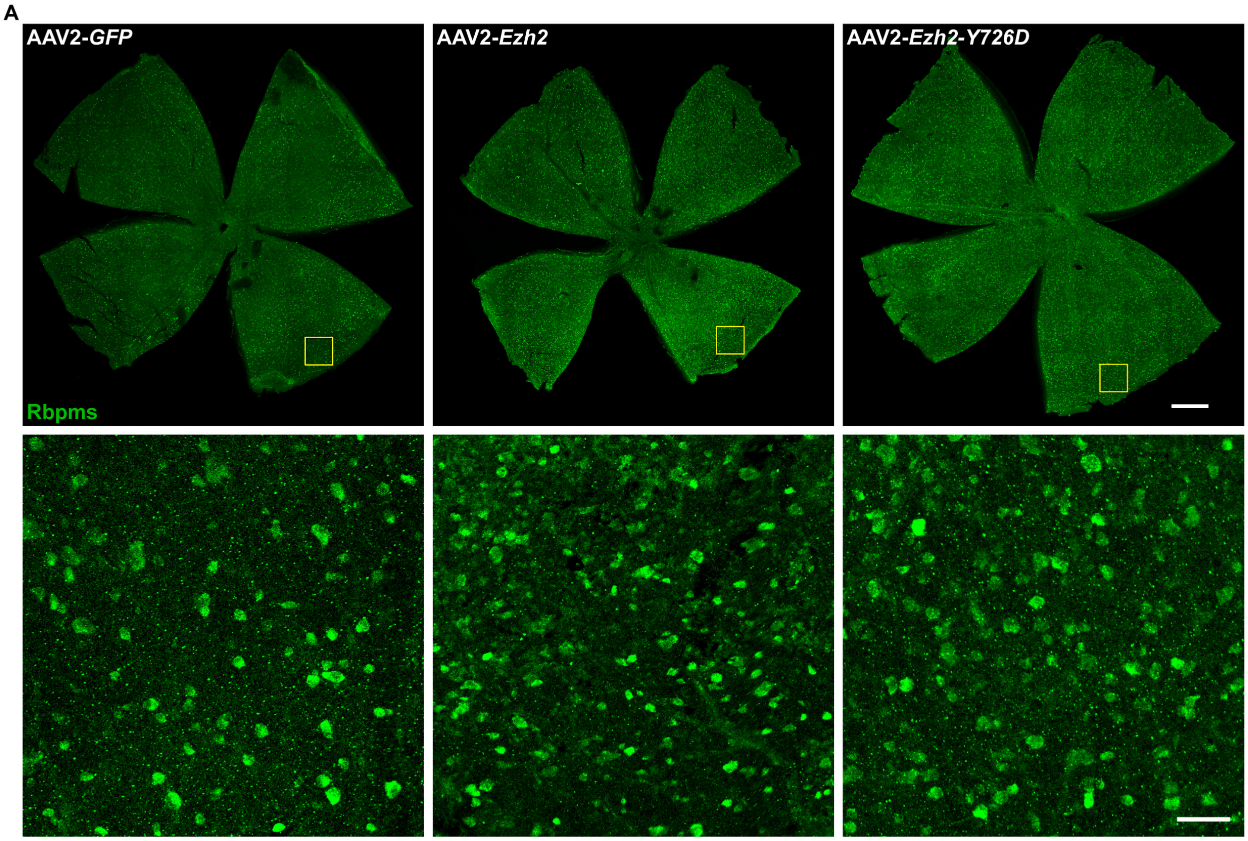


**Figure S2. *Ezh2* overexpression enhances RGC survival after optic nerve crush. Related to Figure 2.**

Immunohistochemistry of whole-mount retinas showing that overexpression of *Ezh2* or *Ezh2-Y726D* improves RGC survival two weeks after optic nerve crush. Whole-mount retinas were stained with anti-Rbpms (green). The lower row displays enlarged images of the areas in yellow boxes in the upper row. Scale bar, 500 µm for the upper row, 50 µm for the lower row.

Figure S3

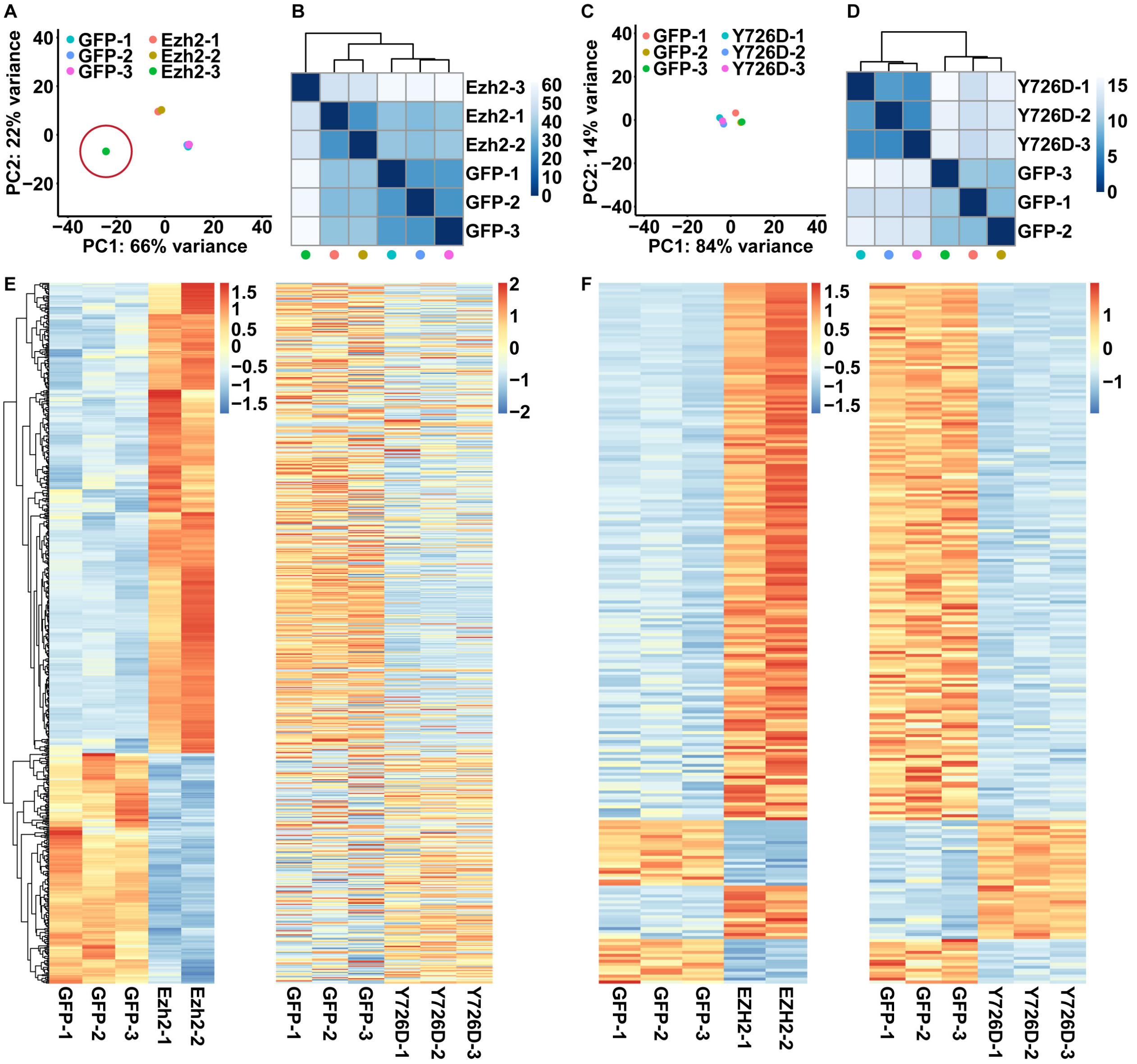
A
GFP-1
GFP-2
GFP-3
Ezh2-1
Ezh2-2
Ezh2-3
PC1: 66% variance
PC2: 22% variance
B
Ezh2-3
Ezh2-1
Ezh2-2
GFP-1
GFP-2
GFP-3
C
GFP-1
GFP-2
GFP-3
Y726D-1
Y726D-2
Y726D-3
PC1: 84% variance
PC2: 14% variance
D
Y726D-1
Y726D-2
Y726D-3
GFP-3
GFP-1
GFP-2
E
GFP-1
GFP-2
GFP-3
Ezh2-1
Ezh2-2
GFP-1
GFP-2
GFP-3
Y726D-1
Y726D-2
Y726D-3
F
GFP-1
GFP-2
GFP-3
EZH2-1
EZH2-2
GFP-1
GFP-2
GFP-3
Y726D-1
Y726D-2
Y726D-3

**Figure S3. Ezh2-Y726D acts as a dominant-negative form of Ezh2 in RGCs. Related to Figure 4.**

(A, B) Principal component analysis (A) and hierarchical clustering (B) of RNA-seq libraries of control and *Ezh2* overexpression conditions. Note that one library in the *Ezh2* overexpression condition (circled in A) was excluded from further analysis due to low repeatability with other two libraries.

(C, D) Principal component analysis (C) and hierarchical clustering (D) of RNA-seq libraries of control and *Ezh2-Y726D* overexpression conditions. Note that the control (GFP) libraries in (C, D) are independent of those in (A, B).

(E) Heatmaps of the 669 differentially expressed genes regulated by *Ezh2* overexpression in the control vs. *Ezh2* overexpression RNA-seq (left) or the control vs. *Ezh2-Y726D* overexpression RNA-seq (right). The same gene is represented in the same row on the left and right.

(F) Heatmaps of the 236 common differentially expressed genes regulated by both Ezh2 overexpression and *Ezh2-Y726D* overexpression in the control vs. *Ezh2* overexpression RNA-seq (left) or the control vs. *Ezh2-Y726D* overexpression RNA-seq (right). The same gene is represented in the same row on the left and right.

Figure S4

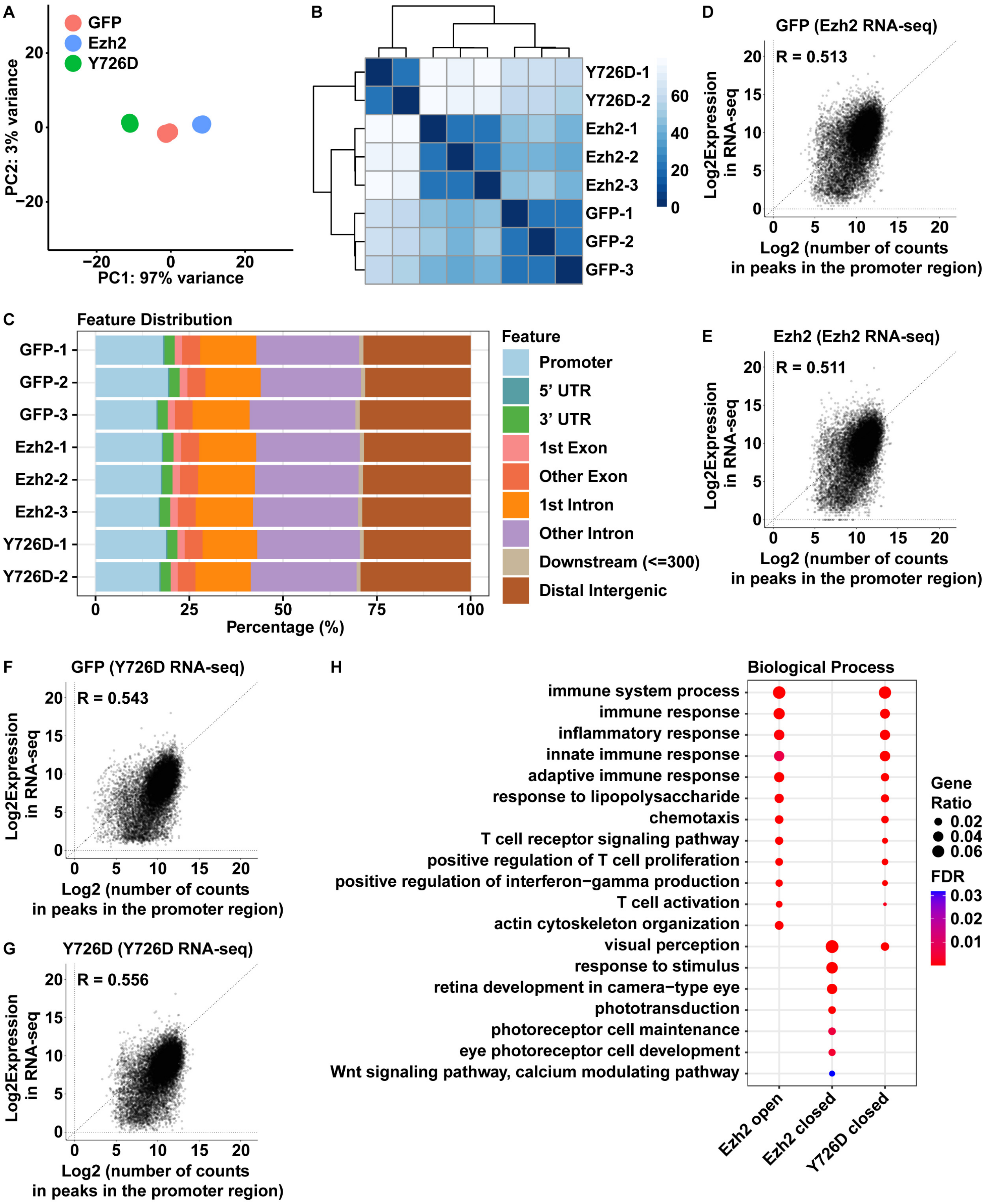
A
GFP
Ezh2
Y726D
PC2: 3% variance
PC1: 97% variance
B
Y726D-1
Y726D-2
Ezh2-1
Ezh2-2
Ezh2-3
GFP-1
GFP-2
GFP-3
D
GFP (Ezh2 RNA-seq)
R = 0.513
Log2Expression in RNA-seq
Log2 (number of counts in peaks in the promoter region)
C
Feature Distribution
GFP-1
GFP-2
GFP-3
Ezh2-1
Ezh2-2
Ezh2-3
Y726D-1
Y726D-2
Percentage (%)
Feature
Promoter
5' UTR
3' UTR
1st Exon
Other Exon
1st Intron
Other Intron
Downstream (<=300)
Distal Intergenic
E
Ezh2 (Ezh2 RNA-seq)
R = 0.511
Log2Expression in RNA-seq
Log2 (number of counts in peaks in the promoter region)
F
GFP (Y726D RNA-seq)
R = 0.543
Log2Expression in RNA-seq
Log2 (number of counts in peaks in the promoter region)
G
Y726D (Y726D RNA-seq)
R = 0.556
Log2Expression in RNA-seq
Log2 (number of counts in peaks in the promoter region)
H
Biological Process
immune system process
immune response
inflammatory response
innate immune response
adaptive immune response
response to lipopolysaccharide
chemotaxis
T cell receptor signaling pathway
positive regulation of T cell proliferation
positive regulation of interferon−gamma production
T cell activation
actin cytoskeleton organization
visual perception
response to stimulus
retina development in camera−type eye
phototransduction
photoreceptor cell maintenance
eye photoreceptor cell development
Wnt signaling pathway, calcium modulating pathway
Gene Ratio
0.02
0.04
0.06
FDR
0.03
0.02
0.01
Ezh2 open
Ezh2 closed
Y726D closed

**Figure S4. ATAC-seq results are consistent with RNA-seq results. Related to Figure 4.**

(A-C) Principal component analysis (A), hierarchical clustering (B), and genomic feature distribution (C) of ATAC-seq libraries of control, *Ezh2* overexpression, and *Ezh2-Y726D* overexpression conditions.

(D-G) Pearson correlation between RNA expression in RNA-seq and chromatin accessibility at the promoter region in ATAC-seq within each condition.

(H) Gene ontology (GO) analysis of genes whose promoter regions became differentially accessible after *Ezh2* or *Ezh2-Y726D* overexpression. A subset of most significantly enriched GO terms in the biological process category are shown here.

Figure S5

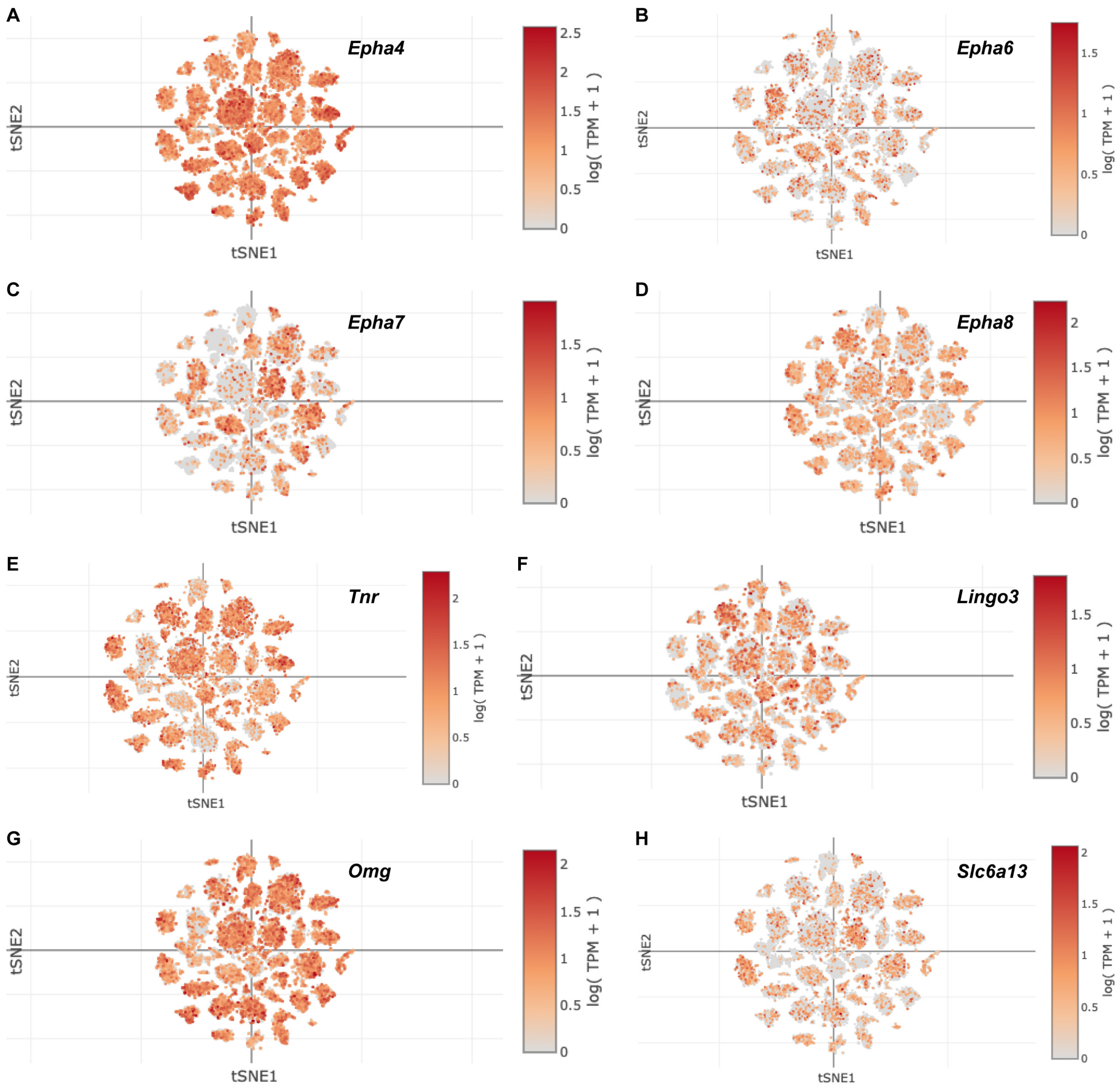


**Figure S5. Expression of a subset of genes regulated by Ezh2 in RGCs. Related to Figure 4-6.**

(A-J) Expression of *Epha4* (A), *Epha6* (B), *Epha7* (C), *Epha8* (D), *Tnr* (E), *Lingo3* (F), *Omg* (G), and *Slc6a13* (H) mRNAs in RGCs shown by t-SNE plots obtained from https://singlecell.broadinstitute.org/single_cell/study/SCP509/mouse-retinal-ganglion-cell-adult-atlas-and-optic-nerve-crush-time-series (Tran *et al.*, 2019).

Figure S6

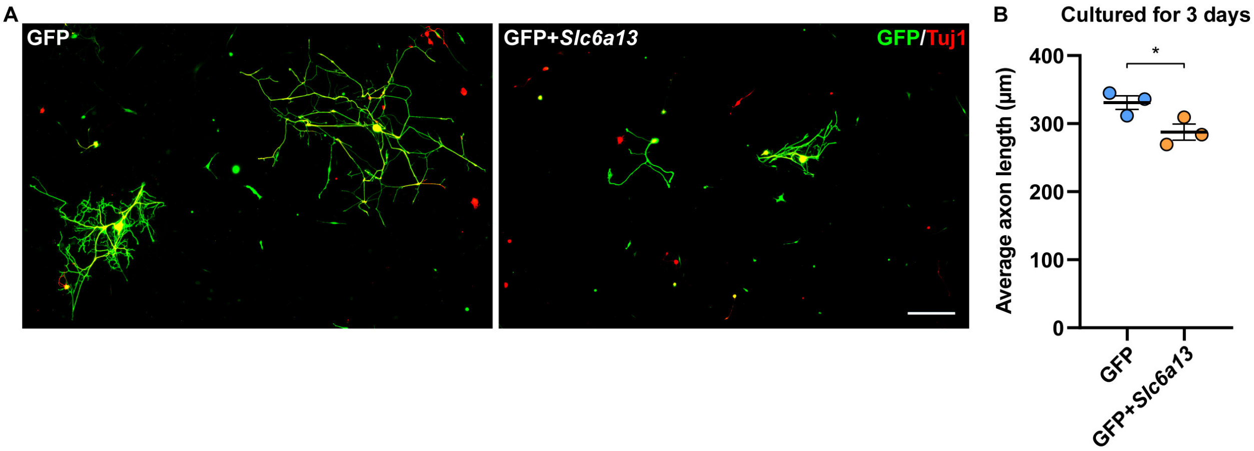


**Figure S6. *Slc6a13* overexpression impairs regenerative axon growth of sensory neurons *in vitro.* Related to Figure 5.**

(A) Representative immunocytochemistry of cultured DRG neurons showing that *Slc6a13* overexpression impairs regenerative axon growth of sensory neurons *in vitro.* Cells were stained with anti-GFP (green) and anti-tubulin β3 (red). Scale bar, 100 μm.

(B) Quantification of the average length of the longest axon of each neuron in (A) (unpaired t test; $p = 0.0481$; $n = 3$ independent experiments; at least 60 neurons were analyzed for each condition in each independent experiment).

Data are represented as mean ± SEM. $^*p < 0.05$.

Figure S7

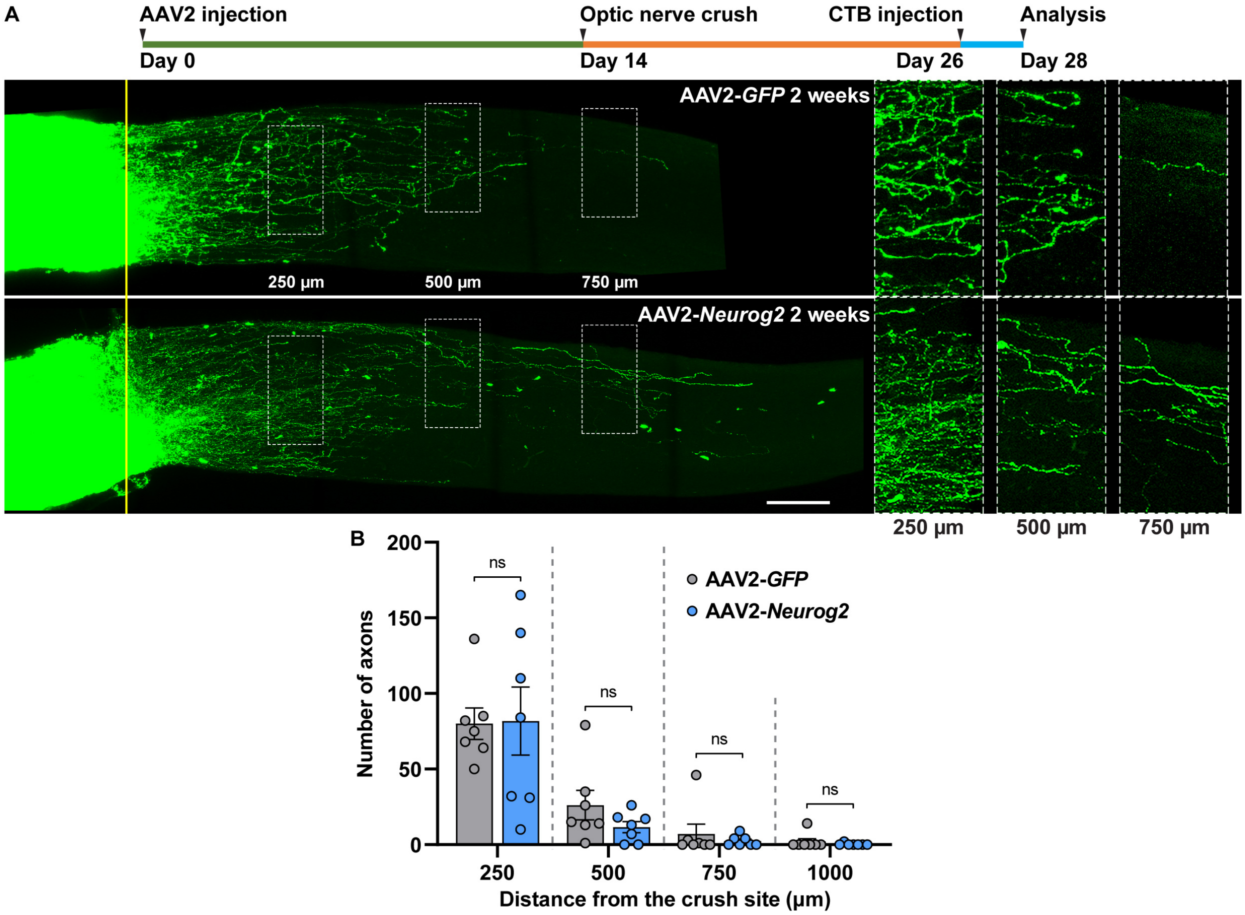


**Figure S7. *Neurog2* overexpression does not promote optic nerve regeneration. Related to Figure 7.**

(A) Top: Time course of the experiment. Bottom: Representative images of optic nerves showing that *Neurog2* overexpression does not promote optic nerve regeneration. Columns on the right display enlarged images of the areas in white, dashed boxes on the left, showing axons at 250, 500, and 750 μm distal to the crush sites, which are aligned with the yellow line. Yellow arrows indicate longest axons in each nerve. Scale bar, 100 μm (50 μm for enlarged images).

(B) Quantification of optic nerve regeneration in (A) (unpaired t test; $p = 0.9460$, 0.1854, 0.4900, and 0.4128 at 250, 500, 750, and 1,000 μm, respectively; $n = 7$ nerves in each condition).

Data are represented as mean ± SEM. ns, not significant.
